# Structural basis of membrane targeting and coatomer assembly by human GBP1

**DOI:** 10.1101/2023.03.28.534355

**Authors:** Tanja Kuhm, Cecilia de Agrela Pinto, Luca Gross, Stefan T. Huber, Clémence Taisne, Evdokia A. Giannopoulou, Els Pardon, Jan Steyaert, Sander J. Tans, Arjen J. Jakobi

## Abstract

Guanylate-Binding Proteins (GBPs) are interferon-inducible guanosine triphosphate hydrolases (GTPases) that mediate immune effector functions against intracellular pathogens. A key step for the antimicrobial activity of GBPs is the formation of homo- and heterooligomeric complexes on the membrane of pathogen-associated compartments or cytosol-invasive bacteria. Similar to other large GTPases of the dynamin family, oligomerisation and membrane association of GBPs depend on their GTPase activity. How nucleotide binding and hydrolysis prime GBPs for membrane targeting and coatomer formation remains unclear. Here, we report the cryo-EM structure of the full-length human GBP1 dimer in its guanine nucleotide-bound state and resolve the molecular ultrastructure of GBP1 coatomer assemblies on liposomes and bacterial lipopolysaccharide membranes. We show how nucleotide binding promotes large-scale conformational changes of the middle and GTPase effector domains that expose the isoprenylated carboxyl-terminus for association with lipid membranes. Our structure reveals how the α-helical stalks of the middle domain form a parallel arrangement firmly held in a unique cross-over arrangement by intermolecular contacts between adjacent monomers. This conformation is critical for GBP1 dimers to assemble into densely packed coatomers on target membranes. The extended α-helix of the effector domain is flexible and permits intercalation into the dense lipopolysaccharide layer on the outer membrane of gram-negative bacterial pathogens. We show that nucleotide-dependent oligomerisation and GTP hydrolysis yield GBP1 membrane scaffolds with contractile abilities that promote the formation of tubular membrane protrusions and membrane fragmentation. Collectively, our data provide a structural and mechanistic framework for interrogating the molecular basis for GBP1 effector functions in intracellular immunity.

## Main

Robust mechanisms for recognition and elimination of microbial pathogens are essential for maintaining the integrity of mammalian organisms. These are the central tasks of the innate immune system and the humoral arm of the adaptive immune system, which cooperate to ensure that a rapid response is formed to eliminate pathogens from extracellular spaces. Yet, many clinically relevant microbes have developed strategies to breach this barrier by surviving and replicating inside host cells (1). As a response, mammalian cells have evolved molecular machinery that elicits effector mechanisms to combat intracellular microbes at the level of individual cells (2). These include pathogen elimination by autophagy, effector immune activation by interferon (IFN) cytokines and the activation of inflammasome complexes (3–7). To subvert cytosolic surveillance, some intracellular pathogens co-opt the host cell endomembrane system to remain sequestered in customised pathogen-associated vacuoles (1, 8). Other pathogens actively disrupt this compartment to replicate in the cytosol (9–11). In either case, the machinery of cell-autonomous immunity forms the last line of defense against such pathogens.

One of the most potent effector systems in cell-autonomous immunity leads to the release of type I and type II interferon cytokines and expression of IFN-stimulated genes. Among the most strongly induced genes is a conserved superfamily of dynamin-like guanosine triphosphatases (GTPases) including the family of Guanylate-Binding Proteins (GBPs) (12, 13). Over the past decade, GBPs have been recognised as key players in mediating host defense against intracellular vacuole-resident and cytosolic bacteria, but also parasites and viruses (14–17). GBPs have been proposed to be recruited to the membrane of some PCVs to form sensory platforms (4, 7), affect vacuolar integrity (6, 18), or to engage with the membrane of cytosolic gram-negative bacteria and parasites directly (16, 18–21).

The human genome encodes seven GBP paralogs sharing similarities with other members of the dynamin-like GTPase superfamily that undergo guanine nucleotide-dependent oligomerisation and mediate diverse biological functions in promoting membrane fusion or fission (22). GBPs have a high intrinsic GTPase activity for hydrolysis of guanosine-5’-triphosphate (GTP) to guanosine-5’-diphosphate (GDP) without the requirement for auxiliary GTPase activating proteins or guanine nucleotide exchange factors (23–25). The enzymology of GBPs is unique among the dynamin superfamily in that some GBPs can also bind GDP with high affinity to produce guanosine-5’-monophosphate (GMP) (26, 27), which can affect bacterial growth and inflammamatory signaling (28). In the absence of infection, GBPs mainly localise to the cytosol, and some associate with endogenous membranes (29). Upon interferon induction, GBPs are recruited to and rapidly assemble into supramolecular coatomers on pathogen-enclosing compartments (PCVs) (18). Recent studies showed that GBPs can also encapsulate cytosolic gram-negative bacteria. In this case, co-recruitment of other effectors and release of lipopolysaccharides (LPS), a glycosylated lipid component of the outer membrane of gram-negative bacteria, activate the non-canonical inflammasome pathway leading to caspase-4 dependent cleavage of gasdermin D and pyroptosis (6, 20, 21, 30). GTPase activity-dependent GBP recruitment to endogenous, PCV or microbial membranes relies on post-translational modifications of a CaaX isoprenylation motif at the carboxyl-terminus. Three members of the human GBP family contain CaaX motifs that lead to covalent attachment of 20-carbon geranylgeranyl (GBP2 and GBP5) or 15-carbon farnesyl moiety (GBP1) and mediate membrane association *in vivo* (29, 31). In its nucleotide-free resting state the farnesyl moiety of GBP1 is buried in a hydrophobic pocket and requires GTP binding to be released (32). The associated conformational changes promote accessibility of its farnesyl anchor for engagement with lipid membranes and can mediate self-oligomerisation into micellar structures in the absence of lipids (33, 34). All reported antimicrobial functions of GBPs are critically dependent on GBP1 isoprenylation, rendering mechanistic insight into the conformational changes that facilitate physical engagement with membranes important for our understanding of their role in cytosolic host defense. In the absence of high-resolution structural data on full-length GBP1 in its activated state, or of native state structures of membrane-associated GBP assemblies, important mechanistic questions related to their mode of action remain unclear.

Here, we determined the cryo-EM structure of the full-length nucleotide-bound dimer of human GBP1. Our structure reveals large scale conformational changes of the α-helical middle domains and GTPase effector domains that stabilise the GBP1 dimer in an extended conformation suitable for association with biological membranes. In vitro biochemical analysis and electron tomography of membrane-assembled farnesylated GBP1 oligomers suggest a critical role of this conformation in GBP coatomer formation on endogenous and bacterial membranes. Importantly, we show that membrane-associated GBP1 assemblies possess GTPase-dependent membrane remodelling capacity that may underlie observations reporting GBP-dependent modulation of membrane integrity and LPS release. Our data provide a structural framework for further studies aiming at unravelling the molecular mechanism of antimicrobial and antiviral activities of GBPs.

## Results

### Cryo-EM structure of the GBP1 dimer

GBP1 is a multidomain protein consisting of an N-terminal large GTPase (LG) domain and a C-terminal α-helical region (C-terminal helical domain, CTHD), which can be further subdivided into a middle domain (MD; α7-α11) and an elongated C-terminal GTPase effector domain (GED; α12-α13) [Fig. 1a] (35). In the absence of guanine nucleotides, the GED folds back onto the MD to span the entire GBP1 molecule and makes extensive interactions with the LG domain and the MD, which are important to maintain GBP1 in the resting state. Quantitative Förster resonance energy transfer (FRET) experiments previously predicted that nucleotide binding and hydrolysis induce major rearrangements of the α-helical stalk region by liberating a latch mediating interaction of α12 with the LG domain (36). This rearrangement results in an extended conformation that releases the C-terminal C15-farnesyl moiety required to reversibly associate with membranes. To map these conformational changes on a structural level we used cryogenic electron microscopy (cryo-EM) to solve the structure of human GBP1 bound to GDP*·*AlF_3_, a compound assumed to represent the transition state of GTP hydrolysis (37). Unlike the isolated LG domain that readily dimerises under several guanine nucleotide conditions (37), full-length GBP1 forms stable dimers only in the presence of GDP*·*AlF_3_ [Fig. 1b,c, Extended Date Fig. S1]. 2D class averages of the GDP*·*AlF_3_-stabilised GBP1 dimer showed one predominant class [Fig. 1d, top panel] making up 92 percent of the total particle set. Only a small subset of classes showed signatures of densities that we assigned to the α-helical stalk [Fig. 1d, middle and lower panel]. Consistently, 3D reconstructions from this data set converged on the LG domain dimer, with no visible density for the stalks comprising MD and GED, and an angular distribution of particles that was indicative of strong preferred orientation [Extended Data Fig. S2]. We hypothesised that the stalks are either highly flexible, or engage in preferential interactions at the air-water interface. To mitigate these effects, we sought to identify strategies to limit the structural heterogeneity by stabilising the extended conformation.

**Fig. 1.**
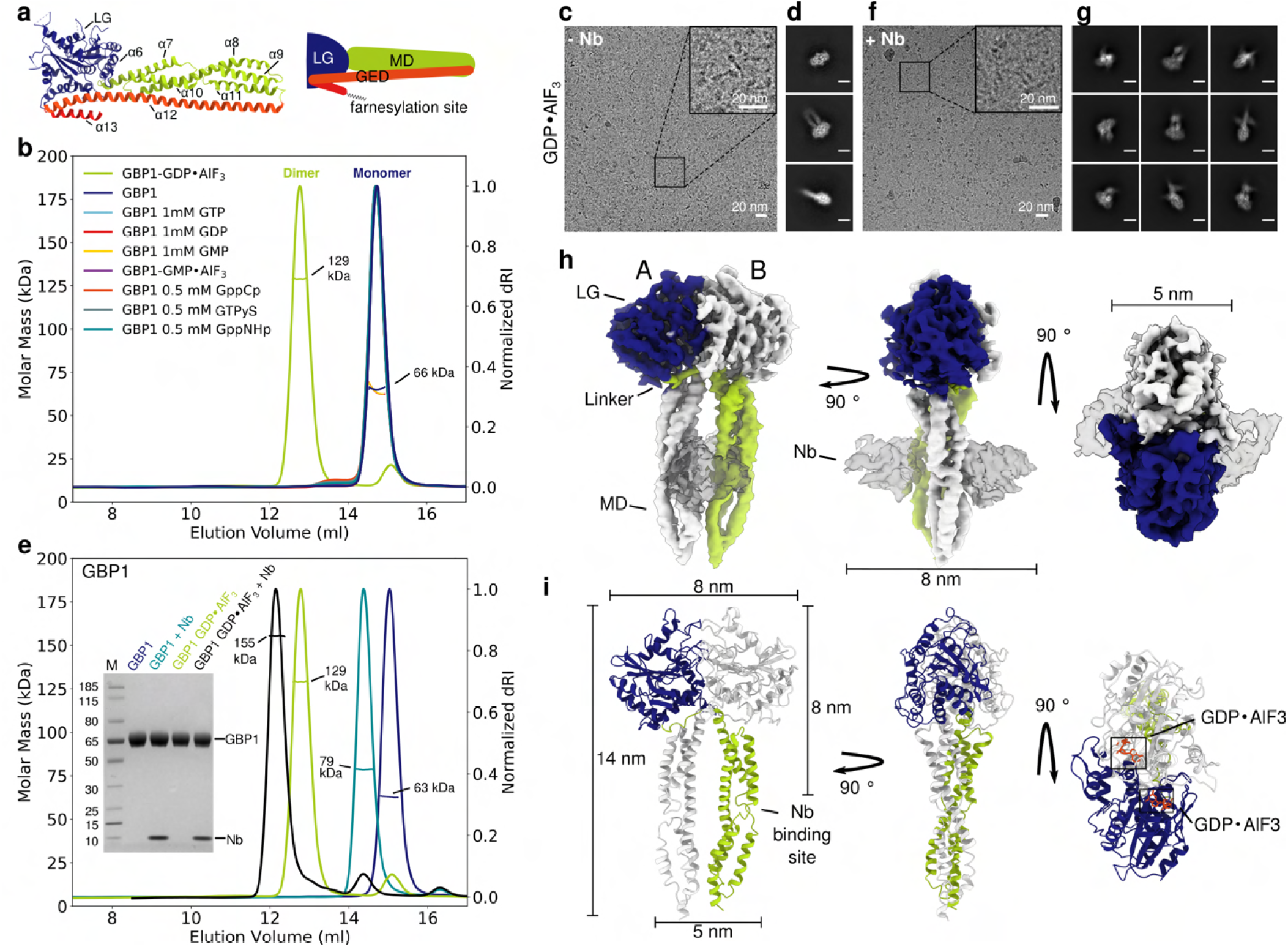
Cryo-EM structure of the GDP*·*AlF_3_ stabilised GBP1 dimer. (a) Atomic model of monomeric GBP1 (PDB ID 1dg3) in cartoon representation alongside a schematic representation displaying the domain architecture. LG: Large GTPase domain (blue), MD: Middle domain (green), GED: GTPase effector domain (orange and red). Individual α-helices in the MD and GED are numbered sequentially. (b) Size-exclusion profile and multi-angle laser light scattering (SEC-MALS) analysis of GBP1 with different nucleotides. GBP1 appears monomeric on SEC-MALS in the presence of GTP, GDP, GMP, GMP·AlF_3_, GppCp, GTPyS or GppNHp, while a dimer peak emerges in the presence of GDP*·*AlF_3_ . The experimentally determined molecular weight is plotted across the chromatographic peak and is reported in kDa. (c) Representative cryo-EM micrograph and (d) 2D class averages of GBP1-GDP*·*AlF_3_ . Scale bar in (d) is 5 nm. (e) Size-exclusion profile and MALS analysis of recombinant GBP1:Nb74 complex in the presence and absence of GDP*·*AlF_3_ . An SDS-PAGE analysis of peak fractions is also shown (molecular marker in leftmost lane, in kDa). (f) Cryo-EM micrograph and (g) 2D class averages of GBP1-GDP*·*AlF_3_ -Nb74 (scale bar 5 nm). (h) 3D cryo-EM density map of the GDP*·*AlF_3_ -stabilised GBP1-dimer bound to Nb74. (i) Refined atomic model of the GBP1-dimer as derived from fitting into the cryo-EM density in (h). The nucleotide binding sites located at the LG dimer interface are highlighted in orange.

### A novel nanobody specific for the GBP1 middle domain

We raised camelid antibodies (nanobodies) specific for human GBP1 by immunising llamas with recombinant GBP1. We then constructed a phage display library from mRNA isolated from peripheral lymphocytes (38) and selected nanobodies specific for GBP1 by panning, which were confirmed by ELISA using immobilised GBP1. We next tested a subset of nanobodies from different complementarity-determining region (CDR) clusters on their ability to bind the GDP*·*AlF_3_-stabilised GBP1 dimer using SEC-MALS and mass photometry. Of several candidates, we selected one nanobody (Nb74) that bound both GBP1 monomers and dimers in an apparent 1:1 molar ratio [Fig. 1e, Extended Data Fig. S3, Extended Data Table S1]. To confirm whether Nb74 binds the extended stalk, we acquired cryo-EM micrographs of GDP*·*AlF_3_ stabilised GBP1 dimer in the presence of Nb74. 2D class averages revealed pronounced density protruding from the LG dimer, suggestive of better preservation of the α-helical stalk in this sample. Several 2D classes also showed additional density in close proximity to the MD, indicating that Nb74 selectively binds the α-helical region in the extended conformation [Fig. 1f-g].

### Large conformational changes induce a cross-over arrangement of the MDs in nucleotide-bound GBP1

We next determined the 3D structure of Nb74-bound GBP1 dimers in complex with GDP*·*AlF_3_, yielding a pseudo-C2 symmetric 3D reconstruction at a nominal resolution of 3.7 Å [Fig. 1h-i, Extended Data Fig. S4]. We found better resolved density in the stalk region when not imposing C2 symmetry, suggesting some residual structural flexibility of the MD and GED in the dimer which was supported by local resolution analysis and flexible refinement [Extended Data Fig. S4, S5 and Extended Data Movie SM1].

GBP1 associates into dimers via the LG domains, with additional contacts formed between the MDs. The Nb74 nanobody binds the MD at the junction formed by helices α7-8 and α10-11 [Fig. 1h]. In the dimer the MDs form a cross-over arrangement in which the linker regions Gly^307^-Val^316^ connecting LG domain and MD cross each other, such that the MDs associate with the LG domain of the respective pairing monomer and extend in parallel from the LG dimer. The GED (residues 481-592) is likely flexible and not visible in our structure. The LG dimer interface is stabilised by a large contact surface formed across the A-face of the GTPase domain, induced by cross-monomer coordination of the GDP*·*AlF_3_ ligand [Extended Data Fig. S6] and is consistent with crystal structures of the nucleotide-bound LG domain dimer (37). The conformational change induced by nucleotide binding leads to partial unraveling of the C-terminal part of helix α6 (Ile^304^-Ser^306^) and the N-terminal end of helix α7 (Cys^310^ - Val^316^) to form the linker region of the domain cross-over in which the MD swings out to form a new interface with the respective pairing monomer in the GBP1 dimer [Fig. 2a-d]. This interface is primarily formed by a hydrophobic patch between the LG domain of one monomer and an aliphatic stretch of residues at the C-terminal end of the linker region and helix α7 (residues Ala^315^ - Ile^322^) in the other monomer [Fig. 2c,d]. While the MDs undergo a large spatial transformation relative to the resting state, their overall conformation of helices α7-α11 remains essentially unchanged with a root mean square deviation (r.m.s.d) of 2.03 Å relative to the nucleotide-free state after superimposing Cα atoms of the MD (residues 327-481). The parallel arrangement of the MD is stabilised by a series of charged residues along helices α9-α11 that form an electrostatic zipper across the protruding stalks [Extended Data Fig. S7]. Towards the C-terminal end of the MD, helices α11 of both monomers come into close proximity and form an additional contact site. While the EM density at this location was not of sufficient quality to identify individual interactions contributing to this interface, it appears to provide additional stabilisation to the pseudo-symmetric parallel MD arrangement. This interpretation is supported by 3D flexible refinement [Fig. 2e, Extended Data movie SM1], which shows that both MDs undergo concerted motion relative to the LG domains. Close inspection of the EM map revealed additional weak density interspersed between helix α3 and α3’ in the LG domain, and protruding beyond the apical end of the MD. To visualise these map regions, we used a model-independent implementation of local density sharpening (39, 40) to improve scaling of weak densities relative to the better resolved parts. This allowed tracing the α3-α3’ loop (residues 156-167) [Extended Data Fig. S8a]. In addition, we observed tubular density protruding from helices α11, which likely corresponds to the N-terminal end of the flexible helices α12 of the GEDs [Extended Data Fig. S8a,b].

**Fig. 2.**
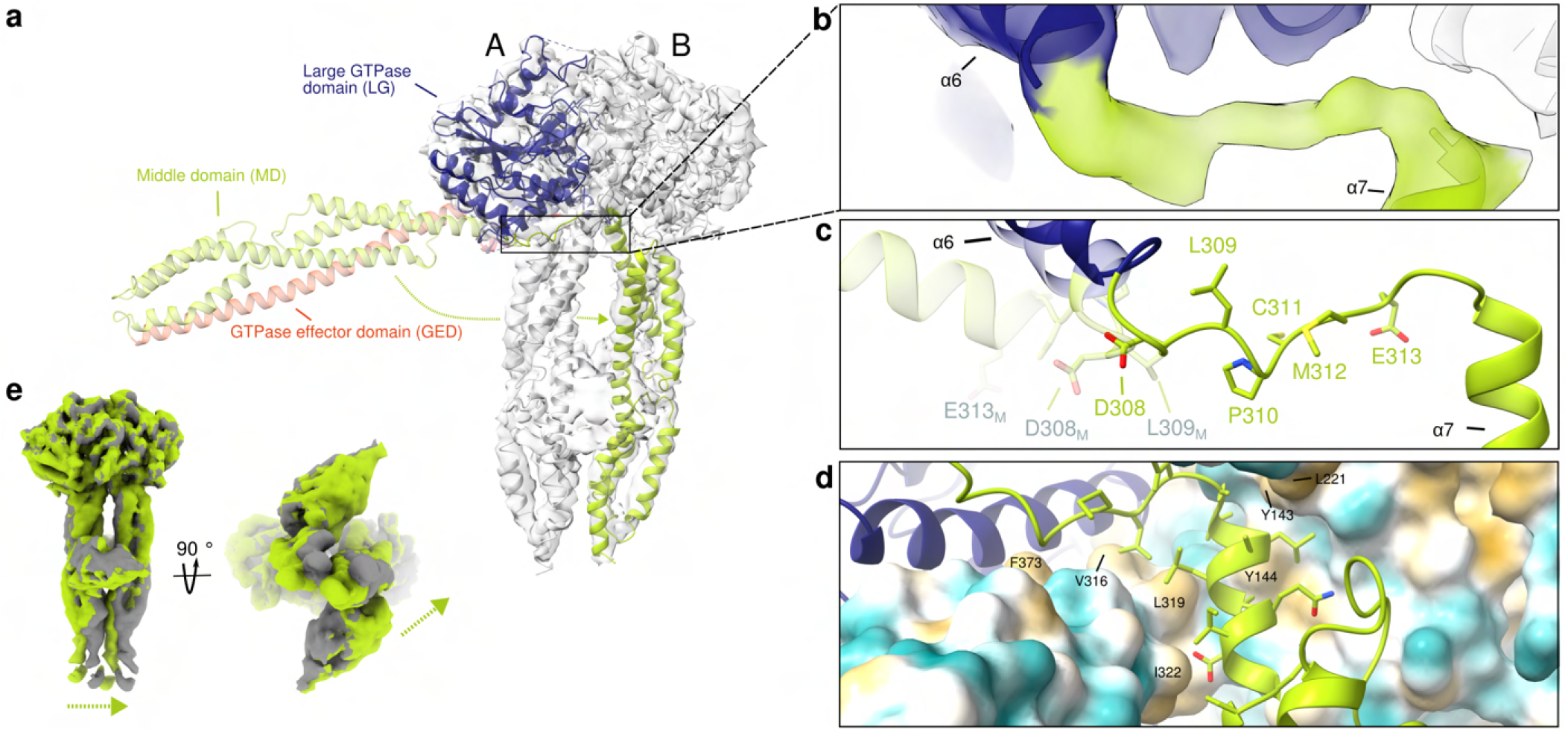
Structural details of the GDP*·*AlF_3_ stabilised GBP1 dimer. (a) The full-length nucleotide-free GBP1 monomer (pale colours; PDB ID 1dg3) is superposed onto one of the monomers in the GDP*·*AlF_3_ stabilised GBP1 dimer (bright colours) by least-squares superposition of their LG domains. Dimer formation involves a large conformational rearrangement in which the α-helical MDs swing across each other to form a parallel arrangement protruding from the LG domains. (b-d) Close-up of the cross-over region (in cartoon representation, superposed on the EM density) formed by a linker region Gly^307^-Val^316^ originating from partially unravelled helices α6 and α7. (c) Residues 307–316 which are part of a linker connecting α6 and α7 need to flip by 180°when changing from the monomer into the dimer conformation. Relevant residues in the nucleotide-free conformation are labelled in grey. (d) The dimer conformation is stabilised by a hydrophobic pocket formed by residues located on α3, α4’ and α7 (highlighted in yellow), interacting with an aliphatic stretch encompassing residues Ala^315^–Ile^322^ on helix α7 of the second monomer. (e) Flexibility of the α-helical MD, visualised through flexible refinement. End points of the density morph along one exemplary latent space dimension are displayed (green and gray densities). The arrow indicates the direction of movement.

### Farnesylated GBP1 forms micellar structures upon nucleotide binding

Our structure of the GDP*·*AlF_3_-stabilised dimer was obtained with recombinant protein devoid of the farnesyl modification at the C-terminal CTIS motif normally required for association with membranes (41). In our structure, the MDs point away from the LG domains as parallel stalks, compatible with a model in which nucleotide binding primes both farnesyl anchors to insert into the membrane. To test this hypothesis, we co-expressed GBP1 together with the human farnesyl transferase machinery to purify natively farnesylated GBP1 (GBP1*_F_*). Unexpectedly, size-exclusion chromatography of GBP1*_F_* in the presence of GDP*·*AlF_3_ did not yield GBP1 dimers as observed for unmodified GBP1 [Fig. 1b]. Instead, in a subset of our SEC-MALS experiments we noted an additional peak corresponding to higher molecular weight species [Extended Data Fig. S3]. Negative stain imaging of this peak fraction revealed circular particles of 58 nm diameter, reminiscent of flowers consisting of discernible petals with a dense spherical perimeter and spoke-like protrusions towards the particle center [Fig. 3a,b]. These structures are consistent with previous observations (33) and were present in higher occurrences if no size-exclusion separation was performed before the imaging experiments. To test whether these structures are exclusively formed with transition-state stabilised GBP1 or more generally occur with actively hydrolysing GBP1, we also prepared samples in the presence of GTP and observed equivalent particles [Fig. 3b] albeit at lower occurrence and requiring higher GBP1*_F_* concentrations. To gain more insight into the molecular architecture of these homo-oligomeric assemblies, we prepared cryo-EM samples of GBP1*_F_* in the presence of GDP*·*AlF_3_ [Extended Data Fig. S9] and performed 2D class averaging [Fig. 3a,c]. Additional electron cryotomograms of GBP1*_F_* micelles revealed that these are 3D micelles and not 2D disc-like structures [Extended Data Fig. S10, Extended Data Movie SM3]. The GBP1 assemblies in these tomograms and the 2D averages appear highly ordered, with approximately 15-16 subunits contained in a semicircle. The increased detail of cryo-EM micrographs of individual flower-like particles allowed us to discern spherical densities at the particle periphery connected to spokes that extend radially towards the center. Based on their dimensions, we ascribe the spherical densities (4.5 nm *±* 0.7 nm in diameter, n = 50) to the LG domains and the spokes to the α-helical stalks. We next quantified the radial dimension of the observed structures to correlate the observed features to our high-resolution model of the GBP1-dimer. If two oppositely oriented GBP1 dimers assemble through interactions at the C-terminus of their respective MDs, the resulting assembly is expected to span 28 nm [see Figure 1i]. Instead, we found the rim-to-rim diameter of the assemblies to be 58.1 nm *±* 1.2 nm (n = 33), suggesting that additional structural elements need to make up the remaining distance. To map the location of the MD within the flower-like assembly, we incubated GBP1_F_ with GDP*·*AlF_3_ in the presence of Nb74. Negative stain images of this sample contained flower-like particles that contained an additional spherical density at approximately 8 nm radial distance to the peripheral LG domain [Fig. 3d-f]. This estimate is in agreement with the relative distance of Nb74 bound to the MD and the LG domain as determined in our cryo-EM structure of the Nb74-bound GBP1 dimer [Fig. 1h-i]. The overall particle radius of 29 nm is consistent with a GBP1 containing a fully unlatched α12 helix [compare Fig. 1a,i]. This leads us to conclude that the remaining density of the petal towards the particle center must originate from the C-terminal part of the MD and the GED. The particle center in both negative stain (5.6 nm *±* 0.8 nm in diameter (n=27)) and cryo-EM micrographs (6.3 nm *±* 1.1 nm in diameter (n=22)) displays higher contrast than the peripheral LG domain and MD/GED spokes of the petal. Since the diameter of the particles is incompatible with a fully extended conformation of the entire GED, we hypothesise that this density corresponds to a cluster of α13 helices of the GED and the exposed farnesyl anchors [Fig. 3g].

**Fig. 3.**
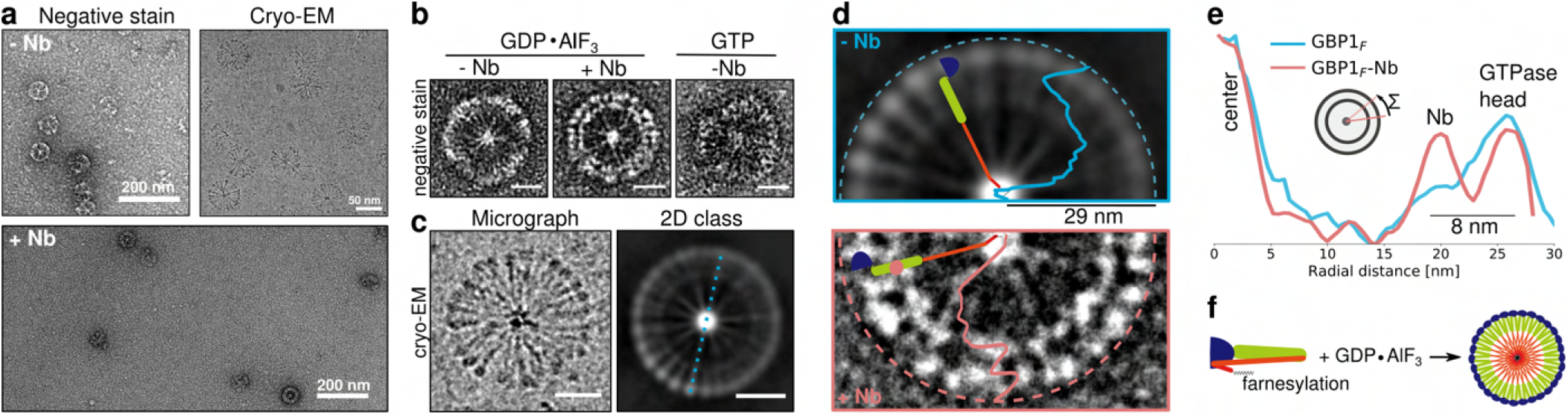
Micellar self-assembly by GBP1*_F_*-GDP*·*AlF_3_. (a) Negative stain images (top left) and cryo EM images (top right) of GBP1*_F_* -GDP*·*AlF_3_ as well as negative stain images of GBP1*_F_* -GDP*·*AlF_3_ with Nb74 (bottom) showing abundant formation of flower-like GBP1 micelles. (b) Close-up of individual GBP1*_F_* -GDP*·*AlF_3_ micelles formed by GBP1*_F_* alone or in the presence of Nb74 (GBP1*_F_* :Nb74). A GBP1*_F_* micelle formed in the presence of GTP is also shown (scale bar 20 nm). (c) Close-up of a cryo-EM micrograph with a GBP1*_F_* -GDP*·*AlF_3_ flower showing discernible repetitive elements (left panel). Cryo-EM 2D class average of GBP1 micelles (right panel) highlighting spherical densities at the perimeter and spokes pointing towards the particle center (scale bar 20 nm). (d-e) Radially averaged intensity profiles of GBP1 micelles plotted on top of a cryo-EM 2D class average of GBP1*_F_* -GDP*·*AlF_3_ micelle (top panel) or on top of a negative stain image of a GBP1*_F_* -GDP*·*AlF_3_ micelle formed with Nb74. A second ring of globular density is visible at 8 nm distance from the perimeter. (f) Schematic representation of GBP1*_F_* -GDP*·*AlF_3_ micelle formation.

### GBP1*_F_* forms dense coatomers on liposomes and scaffolds tubular membrane protrusions

Fluorescently labelled GBP1*_F_* (GBP1*_F_*-Q577C-AF647) uniformly stains brain polar lipid extract (BPLE)-derived giant unilamellar vesicles (GUVs), suggesting homogeneous GBP1 coverage based on fluorescence [Fig. 4a]. To determine at the structural level if the GBP1 conformation observed in the lipid-free GBP1*_F_* micelles is relevant for GBP1 association with membranes, we mixed BPLE-derived small unilamellar vesicles (SUVs) with GBP1*_F_* -GDP*·*AlF_3_ or GBP1*_F_* - GTP for TEM analysis. Cryo-EM micrographs of these samples showed SUVs densely covered with a proteinaceous coat of 29.6 nm (*±* 3.1 nm, n=47) radial extension [Fig. 4b-d], consistent with the dimensions of the extended GBP1*_F_* conformation observed in the lipid-free GBP1 micelles. Strikingly, we observe either fully coated SUVs or SUVs devoid of any GBP1 coat [Fig. 4b,c], suggesting that cooperativity in membrane association may affect efficacy of GBP1 coating potential. On a subset of SUVs, we observed extended tubular protrusions of 59.8 nm (*±* 2.4 nm, n=37) diameter scaffolded by GBP1 in an arrangement indistinguishable from that on spherical liposomes at the level of detail discernible from these images. Cryo-EM micrographs of such structures in unsupported ice revealed these protrusions to be highly flexible [Fig. 4e], precluding 2D averaging. Negative stain analysis at different GBP1 concentrations revealed that the formation of protrusions was highly concentration dependent, transitioning from uniformly coated SUVs through buds that buckle outwards and finally to scaffolded tubule extrusion beyond a certain threshold concentration [Fig. 4f]. 2D class averages and associated power spectra of negatively stained protrusions show repetitive features that are consistent with overall dimensions of laterally associated GBP1 molecules [Extended Fig. S11, Fig. 1i]. The micrographs did not allow us to uniquely discriminate whether the protrusions contain membrane or are formed by excess GBP1 through aggregation of exposed farnesyl anchors similar to GBP1 micelle formation. To test these possibilities we also performed concentration series experiments with GBP1*_F_*in the absence of lipids. Under these conditions, we observed increased formation of micellar assemblies but no formation of tubular structures, suggesting that tubule formation by GBP1 involves extrusion of membrane material [Extended Data Fig. S12].

**Fig. 4.**
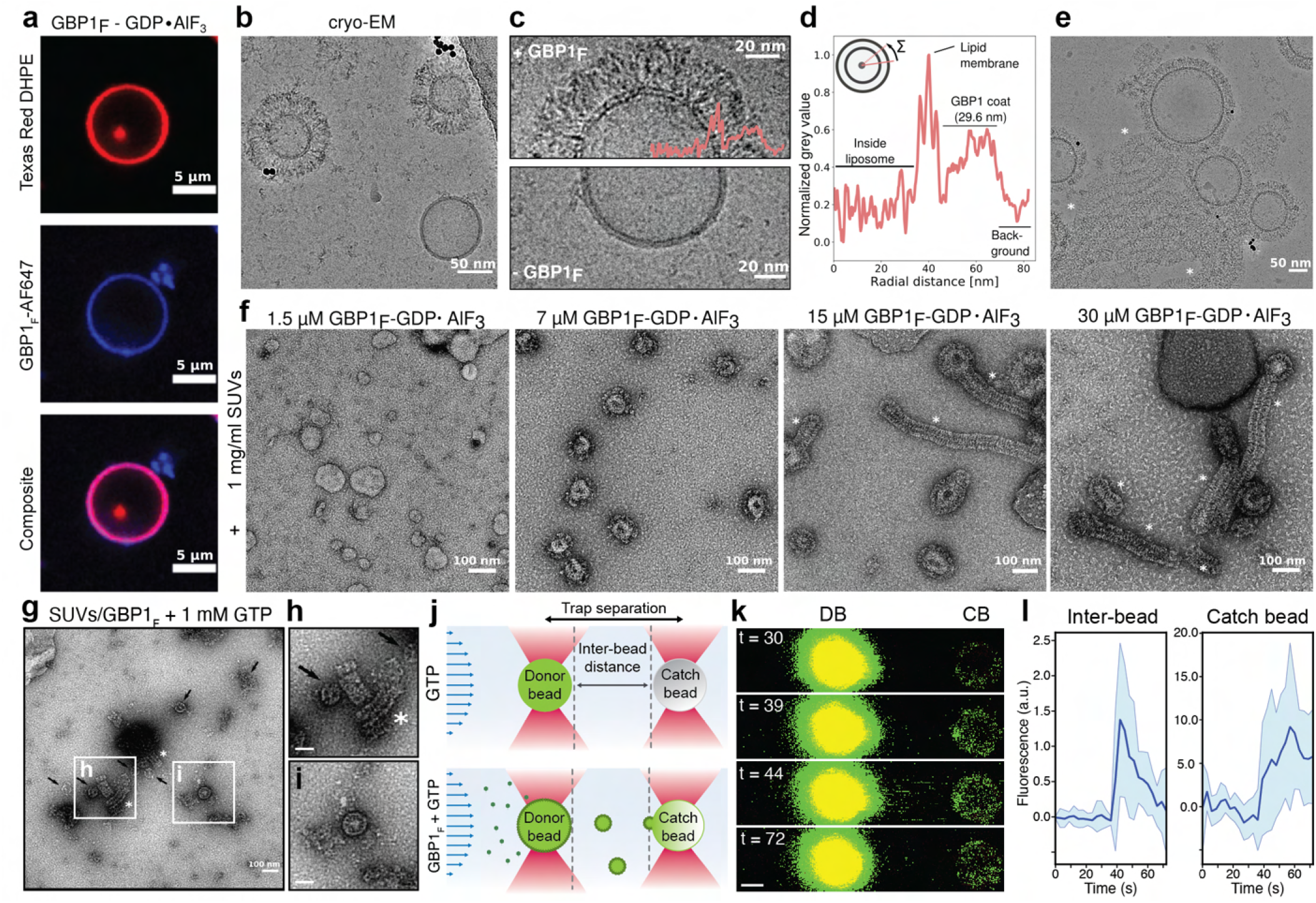
Coatomer formation and membrane remodeling capability of GBP1. (a) Confocal fluorescence imaging of Alexa647-labelled GBP1*_F_* binding to DHPE-texas red-labelled BPLE GUVs in the presence of GDP*·*AlF_3_ . (b) Cryo-EM micrograph of GBP1*_F_* -GDP*·*AlF_3_ binding to BPLE liposomes. (c) Close-up of BPLE SUVs with or without GBP1*_F_* -GDP*·*AlF_3_ coat. (d) Radially averaged intensity profile across a GBP1*_F_* -coated SUV (e) Cryo-EM micrographs of tubular protrusions (asterisks) formed on GBP1-coated SUVs. (f) GBP1 coatomer formation is concentration dependent. A visible coat starts to form at GBP1*_F_* concentrations of 7 µM. At a concentration of 15 µM and higher, GBP1-coated tubular protrusions (asterisk) become visible and are the dominant structures at concentrations exceeding 30 µM. (g-i) In the presence of GTP, GBP1*_F_* remodels SUVs into spherical micelles (arrows) and short filaments (asterisks). Scale bar in (h-i): 50 nm. (j) Dual-trap membrane fragmentation assay: A donor bead coated with rhodamine 6G-labeled membrane is held in place by an optical trap in a flow cell operating at constant pressure. A second, uncoated “catch” bead is held in an additional optical trap at 6 µm distance of the donor bead. Lower panel: GBP1*_F_* -dependent membrane scission/fragmentation would result in membrane transfer from the bead-supported donor membrane to the catch bead. The schematic transfer of membrane vesicles is shown for illustrative purposes only and the precise structure of membrane fragments and GBP1*_F_* scaffold is unclear. (k) Representative confocal fluorescence images illustrating GBP1*_F_* -dependent lipid transfer from the donor bead (DB) to the catch bead (CB). Scale bar: 1 µm. (l) Integrated fluorescence intensity time traces of the inter-bead space and the catch bead in the presence of GBP1*_F_* and GTP. Solid lines represent mean fluorescence intensities of xx experiments and shaded areas represent 95% confidence intervals.

### GTP hydrolysis promotes GBP1-dependent membrane fragmentation

We next tested whether GBP1 scaffolding of membrane protrusions persists in conditions that allow active GTP turnover. Importantly, for conditions containing GTP we exclusively observe short tubular membrane stubs scaffolded by a GBP1 coat while coated SUVs were absent in our micrographs, suggesting that GTP hydrolysis drives GBP1-dependent scission or fragmentation of liposomes [Fig. 4g-i, S13]. Consistent with the higher GBP1 concentrations required for formation of micelllar assemblies in the absence of lipids, we observed weaker binding to GUVs for equimolar levels of GBP*_F_* in the presence of GTP compared to GDP*·*AlF_3_ [Fig. 4a, Extended Data Fig. S14], providing additional support for a threshold-dependent activity of GBP1. To probe the consequences of GTP-dependent GBP1 coatomer formation on membranes in real time, we made use of a dual-trap optical tweezer assay coupled to confocal fluorescence imaging. Two 2 µm silica beads were held in optical traps at 6 µm trap separation within a laminar flow cell operated at constant pressure. One bead coated with a bilayer membrane containing rhodamine 6G-labelled lipids served as a membrane donor, whereas the second uncoated “catch” bead served to sequester lipid material released from the donor bead [Fig. 4j]. We monitored lipid transfer from the donor bead to the catch bead via fluorescence using confocal microscopy [Fig. 4k, Extended Data Movie SM2]. In the absence of GBP1*_F_*, fluorescence in the inter-bead space and on the catch bead remained at constant baseline level [Fig. 4l]. We then dispensed GBP1*_F_* into the flow channel in close proximity to the donor bead and in the presence of GTP. If GTP-dependent membrane scaffolding by GBP1*_F_* results in membrane fragmentation/scission, membrane fragments released from the donor bead will be sequestered by the catch bead under continuous flow. Indeed, we found lipid fluorescence in the inter-bead space and on the catch bead to increase *∼*2- and *∼*7-fold, respectively, approximately 30 seconds after addition of GBP1*_F_* (n=18) [Fig. 4l]. This suggests that lipid material is released from the donor bead in a GBP1*_F_*-dependent manner. Altogether our data indicate that GBP1 scaffolding can promote severing of bilayer membranes and lipid release under conditions that permit GTP hydrolysis.

In tomographic reconstructions of GBP1 assemblies on SUVs we observe that GBP1 covers the entire SUV by assembling into three-dimensional coatomers, which appear to be stabilised by tight lateral association of GBP1 subunits [Fig. 5a-b, Extended Data Movie SM3 and SM4]. The partially regular appearance of the GBP1 coat in individual z-slices from these tomograms is indicative of short range order of the GBP1 coat, and appears to be mediated primarily via interactions of adjacent LG domains. To test this hypothesis, we also acquired cryo-EM micrographs of GBP1-coated SUVs in the presence of Nb74, which through its interactions with the MD domain may sterically affect the lateral association of GBP1 dimers in a dense coatomer [Fig. 1h,i]. Indeed, for these conditions we frequently observed SUVs that were only partially coated and showed signs of structural disorder in the protein coat [Fig. 5c], supporting an important role of an unperturbed GBP1 dimer conformation in coatomer assembly.

**Fig. 5.**
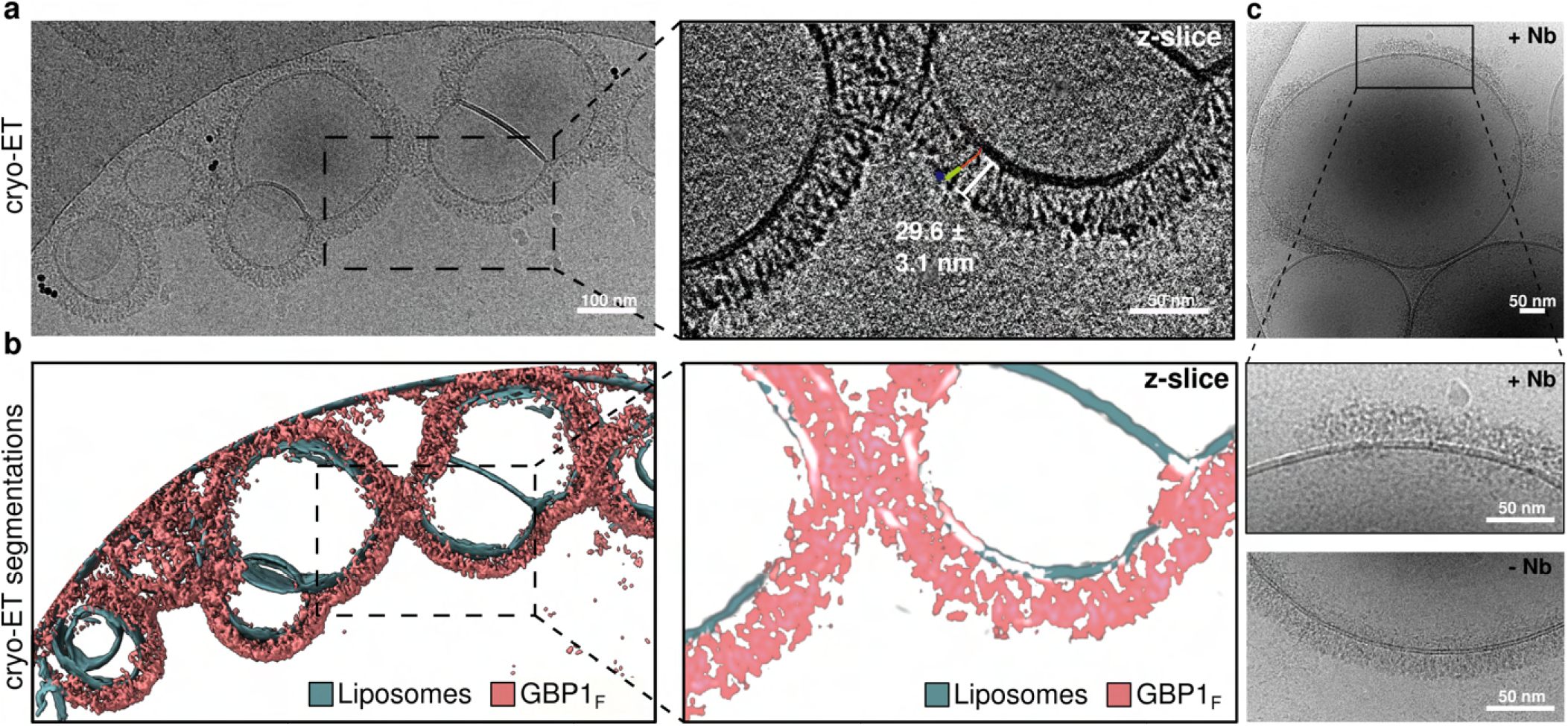
Electron cryo-tomography of GBP1 coatomers. (a) Electron cryo-tomogram of GBP1-coated liposomes. Projected z-stack of reconstructed tomogram and close-up of an individual z-slice showing discernible repetitive subunits. (b) Segmented tomogram from (a); GBP1: pink, membranes: green. (c) Cryo-EM micrograph of GBP1*_F_* - coated liposomes in the presence of Nb74. The close-up shows a comparison to GBP1*_F_* -coated liposomes in the absence of Nb74 (lower panel). The presence of Nb74 results in only partially coated liposomes with a higher degree of structural disorder.

### The cross-over conformation of the GBP1 dimer is critical for membrane association

We next asked whether the cross-over arrangement of the nucleotide-activated GBP1 dimer is required for membrane association. To test this hypothesis, we sought to identify mutants that disrupt the interfaces stabilising this extended conformation but retain the ability to form dimers by association through the LG domains. First, we analysed sequence conservation in the α6-α7 region forming the loop structure in the cross-over conformation, the MD-LG interface, the electrostatic zipper motif and the C-terminal contact site of the MD, and designed point mutants to weaken conserved motifs and interactions [Fig. 6a-b; Extended Data Fig. S15 and S16]. The main interface in the GBP1 dimer is formed between the LG domains of both monomers, contributing 2138 Å^2^ (62%) of the total buried surface area of the dimer interface as inferred from our structure. We therefore expected the different variants to retain their ability to form dimers via the LG domain and to perform GTP hydrolysis [Fig. 6c, Extended Data Fig. S17], but to possibly disrupt the parallel arrangement of the MDs and affect the conformation-dependent ability of GBP1 to associate with membranes. To test this hypothesis, we mixed brain polar lipid extract SUVs with GBP1*_F_* variants activated by GDP*·*AlF_3_ and performed co-sedimentation assays followed by quantitative SDS-PAGE analysis of pellet and supernatant fractions. Of four variants tested, two mutants in the cross-over region showed a 33% (D308S; p<0.05) and 44% (D308S/L309A/P310A; p<0.01) reduction in the membrane-bound fraction compared to the WT control [Fig. 6d-e]. These bulk observations are supported by negative stain imaging of SUVs incubated with the two GBP1 variants [Extended Data Fig. S18], showing substantially decreased coatomer formation on individual SUVs but no complete disruption of binding. For variants affecting the LG-MD and MD-MD interface we found no significant effect [Fig. 6e, Extended Data Fig. S19], suggesting that the ability to form the cross-over arrangement is a determining factor for efficient membrane recruitment of GBP1.

**Fig. 6.**
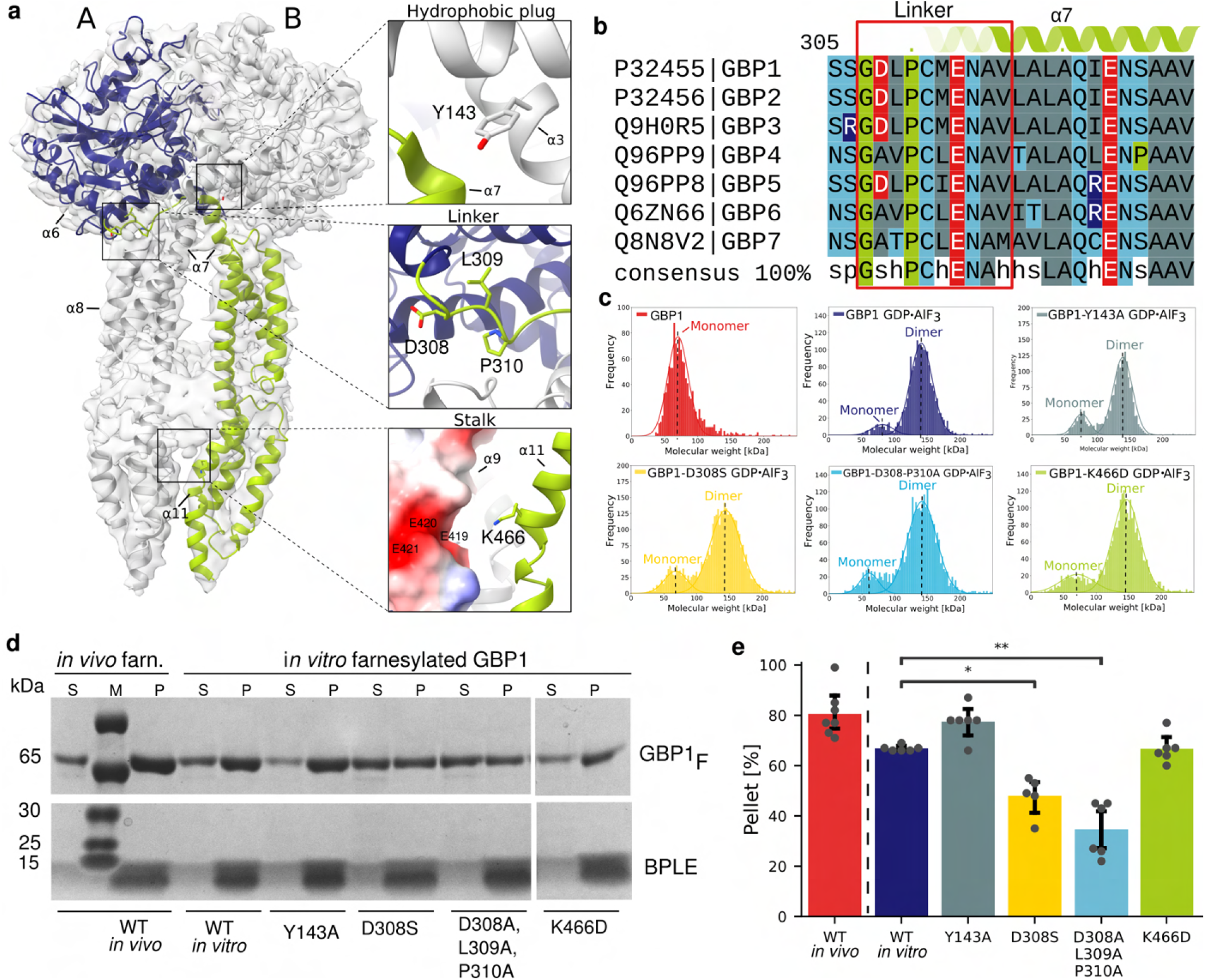
Effect of GBP1 variants on membrane association. (a) Schematic overview of the GBP1 dimer highlighting individual point mutation sites of tested variants. (b) Multiple sequence alignment of all human GBP paralogs zooming in on the linker region (307–316) between α6 and α7. The overall linker region is highly conserved; D308 is conserved across all GBPs containing prenylation motifs (+GBP3). (c) Mass photometry spectra of nucleotide-free GBP1, GBP1-GDP*·*AlF_3_ and GDP*·*AlF_3_ -stabilised GBP1 variants. The determined molecular weights are indicated and show that the point mutations did not affect the ability of the GBP1 variants to dimerise. (d) Representative SDS-PAGE analysis of co-sedimentation assay of GBP1*_F_* variants with BPLE liposomes. S: Supernatant, P: Pellet (e) Quantitative analysis of the co-sedimentation assay. The sum of densitometric intensities of protein in the pellet and supernatant fractions was used to determine the relative percentage of GBP1 in each fraction. The mean intensity and standard deviation are displayed. Statistical significance was determined using Welch’s t-test with Bonferroni correction (*: P≤ 0.05; **: P≤ 0.01)

### GBP1 dimers form the essential unit of GBP coatomers on LPS micelles

Apart from targeting intracellular membranes, GBP1 has been reported to directly associate with lipopolysaccharides (LPS), a glycosylated lipid component of the outer membrane in gram-negative bacteria. LPS consists of a lipid A moiety mediating the integration in the membrane leaflet, a core region of non-repetitive oligosaccharides and the O-antigen consisting of an extended and branched chain of repetitive oligosaccharides. The LPS composition can vary greatly between bacterial strains. To determine whether GBP1 coatomer formation is dependent on the specific oligosaccharide structure of LPS, we incubated nucleotide-activated GBP1*_F_* with three different LPS chemotypes from bacterial pathogens differing in the presence of inner and outer core sugars and O-antigen components; *Salmonella Typhimurium* LPS containing extended O-antigen (LPS-ST), smooth LPS from *Escherichia coli* O111:B4 (LPS-EB) and deep rough LPS from *S. enterica* sv. Minnesota R595 consisting of only the lipid A-Kdo core [Fig. 7a]. Outer-core and O-Ag containing LPS forms elongated bilamellar micelles, whereas deep rough LPS displays semi-vesicular morphology including stretches devoid of a lipid bilayer. For all three chemotypes we observe formation of dense and elongated GBP1 coatomers on remodelled LPS micelles, extending *∼*28 nm from the central lipid bilayer and sandwiching a parallel layer of continuous density with *∼*5-7 nm cross-section, compatible with the estimated thickness of a micellar bilayer including fuzzy contributions of oligosaccharide residues [Fig. 7a]. These dimensions are in agreement with those determined from GBP1 coatomers on brain polar lipid SUVs, suggesting the overall assembly modes of these coatomers are equivalent. As GBP1 forms equivalent coatomers on all LPS forms tested, we conclude that the primary association with LPS membranes is mediated by insertion of the C-terminal farnesyl anchor in the lipid layer, but note that our image data preclude quantitative conclusions on potential affinity differences for certain types of LPS over others. Analogous to micellar GBP1 assemblies and coatomers on lipid SUVs that we use as model system for endogenous membranes, the coat on LPS micelles appears highly ordered with GBP1 molecules assembling in register as concluded from side and top views of coated LPS micelles [Extended Data Fig. S20]. To analyse the detailed mode of GBP1 assembly within the coatomers, we performed 2D class averaging of individual rims of GBP1-coated LPS micelles [Fig. 7b]. 2D class averages of the GBP1 coat revealed low-resolution densities compatible with our high-resolution GBP1 dimer structure viewed in projection, suggesting that GBP1 dimers form the repeating unit in mature coatomers. In some of the classes, we also observed rod-like density extending from the MD towards the LPS lipid surface, supporting a model in which an extended α12 of the GED reaches out for association with the membrane [Fig. 7b]. Together, our results support a model in which nucleotide binding by GBP1 unlatches the C-terminal all-α-helical MD and GED from the LG domain, leading to a swing-like conformational transition of the MD that reassociates with the LG of the adjacent monomer and forms a parallel arrangement of extended GEDs for association with membranes [Fig. 7c]. Interestingly, the dimensions of an extended GED are compatible with the lateral dimensions of extended bacterial LPS O-antigen (LPS-ST: 10.3 *±* 3 nm, (n=23), LPS-EB: 13.1 *±* 2.1 nm (n=11), both measured in negative stain-EM), suggesting that these may be a functional adaption to allow intercalation between the dense O-antigen and core oligosaccharide decoration of the LPS layer and coatomer formation on LPS-containing membranes.

**Fig. 7.**
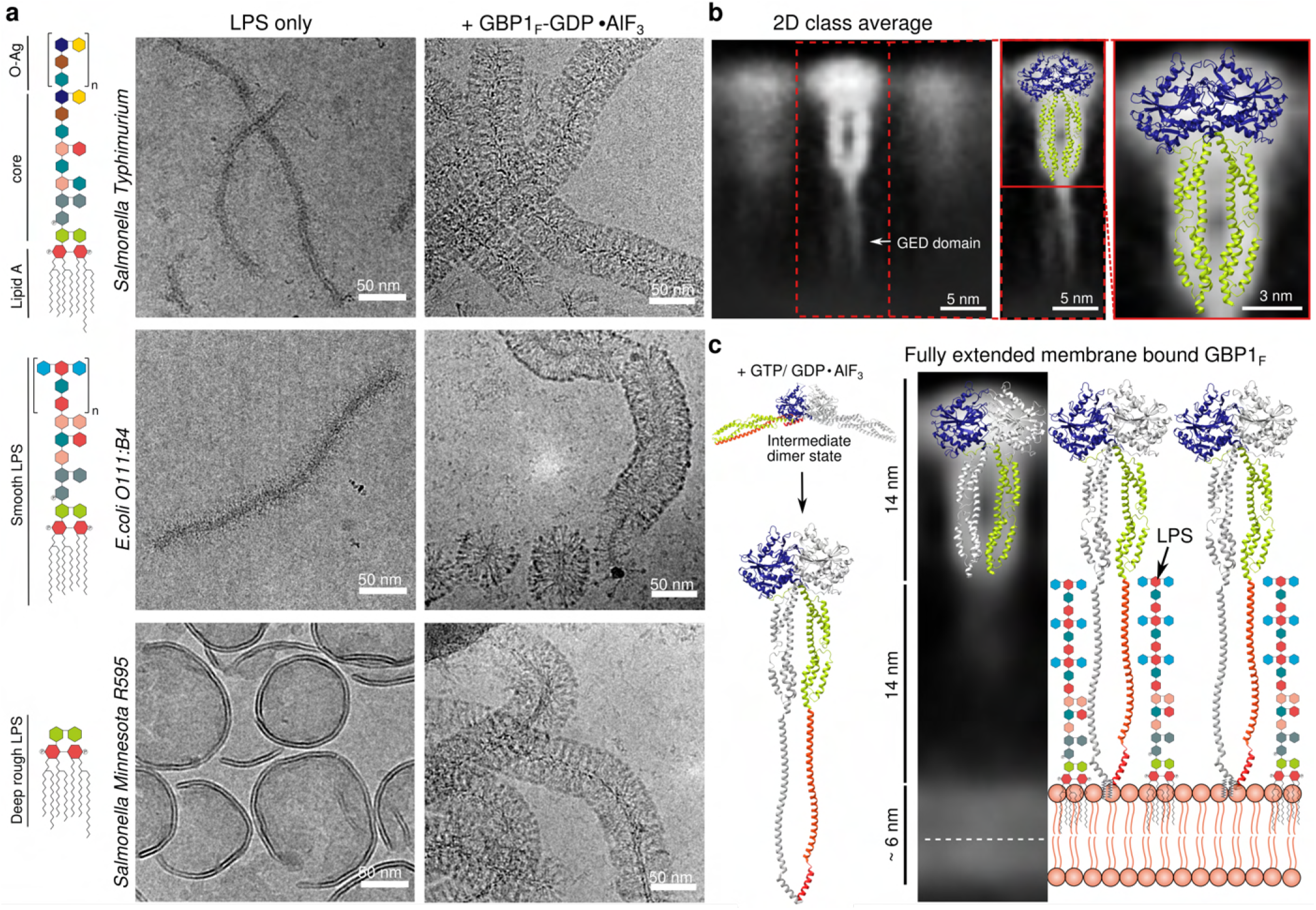
GBP1*_F_* coat formation on LPS micelles of bacterial pathogens. (a) Schematic representation of complex O-antigen containing LPS molecules from *S. enterica* sv. Typhimurium, smooth LPS *E.coli* O111:B4 and deep rough LPS from *S. enterica* sv. Minnesota R595 (green: 2-keto-3-deoxyoctonic acid; grey: L-glycerol-D-manno-heptose; blue-green: galactose; pink: glucose; red: 2-amino-2-deoxyglucose; light blue: Colitose; brown: Rhamnose, yellow: Abequose, darkblue: Mannose). Cryo-EM micrographs of the three types of LPS in the absence (left column) and presence of GBP1*_F_* -GDP*·*AlF_3_ (right column). (b) Selected 2D class average of GBP1*_F_* bound to *E.coli* O111:B4 LPS. Leftmost panels: The extended GEDs projecting toward the membrane surface are visible in the average (white arrow). The atomic model of the GDP*·*AlF_3_ -stabilised GBP1 dimer is superposed onto the projected density. (c) Schematic of nucleotide-dependent activation of GBP1 for membrane binding. Hypothetical encounter complex for initial dimerisation (based on PDB ID 1dg3/2b92), formation of the cross-over conformation of GBP1 dimers upon nucleotide binding with extended GED, and 1D model of the GBP1 coatomer on membranes. The radial extension of O-antigen containing LPS is shown for comparison.

## Discussion

GBPs have recently emerged as important effector molecules in cell-autonomous immunity against intracellular bacteria, and GBP1 forms the central organising unit of this cellular response. The main antimicrobial function of GBP1 has been ascribed to its ability to coat the membrane of pathogen-containing compartments or the outer membrane of gram-negative cytosolic bacteria, where it appears to form a multivalent signaling platform for the activation of the non-canonical inflammasome (16, 17, 20, 21). Coat formation is dependent on nucleotide binding and self-assembly of GBP1. While the functional consequences for GBP1 in intracellular immunity have been firmly established by these studies, the mechanistic underpinnings of these functions remain currently unclear.

Our cryo-EM and cryo-ET data show that the ultrastructure of GBP1 coatomers on lipid and LPS membranes consists of ordered arrays of GBP1 dimers with their α-helical MDs and GEDs protruding in parallel towards the membrane surface. The molecular envelope of the repeating unit that we infer from these data is consistent with our high-resolution cryo-EM structure of the full-length GDP*·*AlF_3_-stabilised GBP1 dimer, displaying a cross-over arrangement of the MD and extended GEDs anchored to the membrane. We found membrane association of GBP1 to be critically dependent on the ability to form cross-over dimers. This cross-over conformation GBP1 is consistent with a recent crystallographic structure of a truncated GBP5 dimer (42) and resembles that of atlastins (43), which are related but functionally different members of the dynamin-like protein (DLP) superfamily of large membrane-associated GTPases. Interestingly, GBPs and atlastins appear to share a set of key structural features stabilising this conformation: a conserved linker region that mediates the MD cross-over, an extended hydrophobic interaction region that latches the MD onto the LG domain of the opposing monomer, and a series of weak interactions holding together the C-terminal α-helices of the MD. While the atlastins and other dynamin-like proteins associate with membranes through through specialised C-terminal domains, transmembrane anchors or amphipathic helices, GBPs are unique among the DLPs in the requirement of isoprenylation for membrane binding. Another distinguishing feature of GBPs is the long, extended α-helical effector domain. While the LG domains and MDs of the GBP1 dimer unit appear rigid, the GED appears to exhibit substantial flexibility. Our data provide important clues for these specialisations. Assembling densely packed coatomers on outer membranes of gram-negative bacteria spiked with extended LPS oligosaccharide chains requires elongated flexible elements that can intercalate between the O-antigen of complex LPS cores. Intriguingly, the dimensions of the extended GED are consistent with the estimated length of LPS chains with extended O-antigen (44), suggesting that the isoprenylated GEDs can act as flexible anchors that allow GBP1 to breach the LPS permeability barrier and assemble densely packed coatomers that are stabilised through interactions between the LG domains of neighbouring dimers.

We found GBP1 coat formation to occur in an all-or-none fashion, with the fraction of coated liposomes strongly dependent on GBP1 concentration. This indicates that coatomer formation is a cooperative process, where successful formation of a GBP coat is dependent on a critical threshold concentration. Cooperativity is a hallmark of processes that require a sharp transition in their biological response, for example by digital activation. The antimicrobial function of GBPs is induced through activation of the interferon pathway that massively upregulates expression of interferon-stimulated genes (ISGs). Interestingly, GBPs are among the most strongly induced ISGs, with basal transcription levels elevated up to three orders of magnitude upon interferon induction (12, 45). A thresholded response to self-assembly may provide GBPs with the ability to prevent coating of intracellular membranes at cellular concentrations under homeostatic conditions and to only activate this function in the presence of infection.

Several recent studies linked GBP1 coatomer formation to the activation of the non-canonical inflammasome pathway, involving recruitment of caspase-4 to the GBP coat and induction of inflammatory cell death (pyroptosis) (16, 20, 21). Pyroptosis is dependent on cleavage of gasdermin D by caspase-4, which in turn is activated by binding to the lipid A component of LPS (46, 47). Caspase-4 dependent pyroptosis is abrogated in the absence of GBP1, suggesting that caspase-4 cannot bind lipid A on bacterial outer membranes by itself. How does GBP1 facilitate access to lipid A components? Our data show that high GBP1 concentrations lead to tubulation of lipid membranes and LPS micelles, indicating that GBP1 has membrane remodelling activity. The tip of membrane tubules forms a region of maximum curvature, which could facilitate access to the membrane-embedded acyl chains of lipid A otherwise shielded by the dense oligosaccharide chains of LPS and therefore inaccessible to the ligand-binding CARD domain of capase-4. A recent study reporting tomographic data on GBP1 coatomer suggested the coat consists of GBP1 monomers (48), which is inconsistent with the GBP1 dimer units observed in our cryo-EM and cryo-ET data reported here and with previous biochemical studies that established homodimer formation as a prerequisite for membrane association of GBP1 (34). Interestingly, LPS and GBP1-dependent retrieval of caspase-4 in cellular pull-downs requires GDP-AlFx (21), confirming the functional relevance of the GBP1 dimer conformation in the membrane-bound oligomers under conditions mimicking the transition-state of GTP hydrolysis. Local remodelling of bacterial membranes by GBP1 oligomers may therefore provide platforms for caspase-4 recruitment and activation and reconciles observations displaying discontinuous GBP1-dependent recruitment of caspase-4 on cytosol-invasive Gram-negative bacteria (21).

All structural data in our study has been acquired using GBP1 arrested in an activated, but non-hydrolysing state. In its native cellular environment GBP1 can bind and hydrolyse GTP, likely leading to further structural rearrangements throughout the hydrolysis cycle. This raises the question of the functional consequences that these structural changes impose on GBP1-coated membranes. Unlike for the transition-state stabilised GBP1 coatomers, we do not observe coated liposomes in the presence of GTP. Instead, we observed fragmented GBP1-decorated membranes (short filaments) or flower-like assemblies resembling, but distinctly different from the micellar structure observed for GBP1*_F_* -GDP*·*AlF_3_ in the absence of lipids. While our present data precludes quantitative conclusions, our observations are indicative of the ability of GBP1 to fragment membranes. How the GBP1 coatomer and GTP-dependent conformational changes relate to this property will be important questions for further studies. Importantly, the concentrations required for GBP1 remodelling activity in the presence of GTP were at least 8-fold increased compared to the situation with GBP1-GDP*·*AlF_3_. Since the concentration of “activated” GBP1 in the presence of GTP will always be lower than that for the non-hydrolysable GTP analog GDP*·*AlF_3_ at equimolar concentrations, this observation is consistent with a threshold-dependent response of GBP1 activity.

In summary, our data firmly establish nucleotide-dependent GBP1 dimers as the functional assembling unit for GBP coatomer formation and demonstrate the capability of this coatomer to utilise GTP hydrolysis to remodel and fragment biological membranes. Important questions, for example how the GBP1 coat is stabilised, how GBP1 recruits and integrates non-prenylated GBP family members into the coatomer, and how these heterotypic interactions affect the functionality of the GBP coat remain unsolved.

## AUTHOR CONTRIBUTIONS

TK, CP and EG purified proteins. TK performed fluorescence imaging, biophysical experiments and prepared cryo-EM samples. TK and AJ collected cryo-EM data. TK, SH and AJ processed cryo-EM data; TK and AJ analysed cryo-EM data and built the atomic model. CP and TK performed mutagenesis and co-sedimentation assays. TK and CT performed cryo-EM of LPS-bound GBP1 coatomers. LG performed, and LG and SJ analysed optical trapping experiments. EP and JS initiated nanobody generation. TK and AJ wrote the manuscript; all authors commented on the final draft. AJ conceptualised and supervised the study.

## DATA AVAILABILITY

The refined atomic model of the pseudo-symmetric GBP1 dimer has been deposited in the Protein Data Bank under accession code 8CQB. The primary cryo-EM density and the LocScale map of the pseudo-symmetric GBP1 dimer are available in the Electron Microscopy Data Bank (EMDB) under accession code EMD-16794. Tomogram reconstructions have been deposited at the EMDB under accession codes EMD-16813, EMD-16814 and EMD-16815. Raw micrographs have been deposited in the Electron Microscopy Public Image Archive (EMPIAR) with accession code EMPIAR-11459. Raw tilt series are available on Zenodo 10.5281/zenodo.7740464.

## Methods

### Plasmid construction

#### GBP1

Codon-optimised synthetic DNA encoding human GBP1 (UniProt accession P32455) was cloned into the NcoI/NotI linearised pETM14 vector containing a N-terminal His_6_ tag and 3C cleavage site, yielding pETM14-GBP1.

#### GBP1 variants

Expression vectors containing GBP1 variants were generated from pETM14-GBP1 by quickchange mutagenesis using appropriate oligos [Extended Data Table S5]. Mutations were confirmed by DNA sequencing (Macrogen Europe B.V., Amsterdam, Netherlands).

#### pCDFDuet-FNTA-FNTB

A co-expression vector for farnesyl transferase (FTase) was constructed using the pCDF-Duet1 (Novagen) vector backbone. FNTA inserts were PCR-amplified with AJLO-023 and AJLO-024 from pANT7-FNTA-cGST (DNASU HsCD00630808). To al low subcloning into MCS1 of pCDF-Duet1, BsmBI sites compatible with NcoI and NotI overhangs were inserted at the 5’- and 3’-ends of FNTA. The BsmBI-digested FNTA fragment was cloned into NcoI/NotI digested pCDF-Duet1, yielding pCDF-Duet-FNTA. For cloning of FNTB into MCS2 of pCDF-Duet-FNTA, FNTB was PCR-amplified from pANT7-FNTB-cGST (DNASU HsCD00077919) using AJLO-25 and AJLO-026 [Extended Data Table S5] to create a 5’-NdeI site and a 3’-BsmBI site compatible with XhoI overhang. An internal NdeI site in pANT7-FNTB-cGST was removed by Quickchange mutagenesis with AJLO-027 and AJLO-028. The NdeI/BsmBI-digested FNTB fragment was cloned into NdeI/XhoI-digested pCDF-Duet-FNTA, yielding the FTase co-expression vector pCDF-Duet-FNTA/FNTB.

#### pCDFDuet-His6-FNTA-FNTB

The pJET1.2 constructs containing FNTA or FNTB were obtained by amplification of AJLD0007 or AJLD0022 [Extended Data Table S6] using AJLO-023 - AJLO-026 [Extended Data Table S5], following the manufacturer recommendations. The pCDFDuet-His6FNTA-FNTB (AJLD0063) vector was obtained by Gibson assembly. The DNA fragments originated from AJLD0052, AJLD0053 and AJLV0038 using primers AJLO-076 - AJLO-083, AJLO-092 and AJLO-093 [Extended Data Table S5]. Successful cloning was confirmed at all stages by DNA sequencing (Macrogen Europe B.V., Amsterdam, Netherlands).

### Protein expression and purification

#### GBP1 and GBP1-variants

Proteins were expressed in *E.coli* BL21(DE3) [Extended Data Table S7] using autoinduction in lactose-containing media. Pre-cultures were grown in LB-medium o/n at 37°C. For protein expression, ZYP5052 medium was inoculated at 1/50 (v/v) with pre-culture and cells were grown at 37°C and 180 rpm for 3-4 h before lowering the temperature to 20°C for 15-20 h. Cells were harvested by centrifugation at 4°C and 4000 rpm and the cell pellet was resuspended in lysis buffer (50 mM HEPES pH 7.8, 500 mM NaCl, 0.1% Triton X-100) on ice. The cells were disrupted by three successive freeze-thaw cycles. To digest genomic DNA, 1-10 ug/ml DNA-seI was added and incubated on a rotating wheel for 1-2 hours at 4°C. To separate cell debris, the lysate was centrifuged at 20,000 x g for 40 min at 4°C. The supernatant was applied to TALON (Takara) affinity resin. The bound fraction was washed with 20 column volumes (cv) of wash-buffer (50 mM sodium phosphate, 300 mM NaCl, 10 mM imidazole, pH 7.4) and eluted in the same buffer containing 150 mM imidazole. The eluent was dialysed into 3C cleavage buffer (50 mM HEPES pH 7.4, 150 mM NaCl, 0.5 mM DTT) and incubated with 1:100 mol/mol 3C protease o/n at 4 °C. Following cleavage, the proteins were further purified via size exclusion chromatography using a GE Superdex200 Increase 10/300 GL column (GE Healthcare) in running buffer (50 mM HEPES pH 7.4, 150 mM NaCl, 0.5 mM DTT).

#### Farnesyl transferase

His-FNTA-FNTB was expressed as described before for GBP1. After harvesting, the cell pellet was placed on ice and resuspended in lysis buffer (50 mM HEPES pH 7.8, 150 mM NaCl, 0.1% Triton X-100). A reduced salt concentration of 150 mM was necessary to avoid disassembly of the FNTA/FNTB . After separating the cell debris, the supernatant was applied to Ni-NTA (GE Healthcare) affinity resin. The bound fraction was washed with 20 column volumes (cv) of wash-buffer (50 mM HEPES pH 7.8, 150 mM NaCl, 0.5 mM DTT, 10-

30 mM imidazole) and eluted in the same buffer containing 250 mM imidazole. The proteins were further purified via size exclusion chromatography using a Superdex200 Increase 10/300 GL column (GE Healthcare) in running buffer (50 mM HEPES pH 7.4, 150 mM NaCl, 0.5 mM DTT).

### In vivo farnesylation of GBP1

Co-translational farnesylation of GBP1 was performed essentially as described (41). *E.coli* BL21(DE3) cells were co-transformed with pETM14-GBP1 and pCDFDuet1-FNTA-FNTB plasmids. The expression and initial purification of GBP1 was performed as described for pETM14-GBP1. To separate non-farnesylated and farnesylated GBP1 (GBP1*_F_*), an additional hydrophobic interaction chromatography (HIC) step was performed. Briefly, NH_4_SO_4_ was added to the protein solution in 3C cleavage buffer to a final concentration of 1 M. The solution was bound to a HiTrap Butyl HP column (GE Healthcare), washed with 30 cv of high salt buffer (1.5 M NH_4_SO_4_, 50 mM Tris-HCl pH 8, 2 mM MgCl_2_, 2 mM DTT) before elution over 20 cv with a linear gradient into low salt buffer (50 mM Tris-HCl pH 8, 2 mM MgCl_2_, 2 mM DTT). Fractions containing the GBP1*_F_* were pooled and further purified by size exclusion chromatography on a Superdex200 Increase 10/300 GL column (GE Healthcare) in running buffer (50 mM HEPES pH 7.4, 150 mM NaCl, 0.5 mM DTT).

### In vitro farnesylation of GBP1

In vitro prenylation of GBP1 was adapted from (49). In brief, 5 µM purified GBP1 was incubated with 5 µM FTase for farnesylation and supplemented with 25 µM farnesyl pyrophosphate (FPP) (Cayman) in prenylation buffer (50 mM HEPES pH 7.2, 50 mM NaCl, 5 mM DTT, 5 mM MgCl_2_, 20 µM GDP). The reaction mixtures was incubated for 60 min at room temperature and dialysed o/n at 4°into running buffer.

### Preparation of GDP·AlF_3_-stabilised GBP1 dimers

15 µM of GBP1 was incubated with 200 µM GDP, 10 mM NaF, 300 µM AlCl_3_, 5 mM MgCl_2_ and 1 mM DTT for 10 min at RT.

### Nanobody generation, selection and purification

#### Nanobody generation

Llamas have been immunised either with purified monomeric GBP1, farnesylated GBP1 or GDP*·*AlF_3_-stabilised dimeric GBP1. From each llama a blood sample was taken and the peripheral blood lymphocytes were isolated followed by the purification of RNA and synthesis of cDNA. Nanobody coding sequences were then PCR-amplified and cloned into a phage display library, creating libraries with *>* 10^8^ independent clones.

#### Nanobody selection

For phage display selections, farnesylated, monomeric or GDP*·*AlF_3_-stabilised dimeric GBP1 was solid phase coated in 50 mM HEPES pH 7.4, 150 mM NaCl, 0.5 mM DTT and selections were performed in the same buffer. To detect the presence of GBP1-specific nanobodies, the His-tag was detected by an anti-His monoclonal antibody followed by the addition of an anti-mouse-antibody conjugated to alkaline phosphatase. As a substrate for alkaline phosphatase conjugates, 2 mg/ml of 4-Nitrophenyl phosphate disodium salt hexahydrate (pNPP) was used. In total 78 clones were found positive on the dimeric GBP1-GDP*·*AlF_3_, 26 on GBP1*_F_* and 33 clones on monomeric GBP1. We selected Nanobodies from different families and performed a SEC-MALS analysis to investigate the binding behaviour. Nanobody 74 was chosen because it binds to GBP1 in a 1:1 ratio without breaking the GDP*·*AlF_3_-stabilised GBP1 dimer apart.

#### Expression and purification

Nanobody 74 (Nb74) was expressed in *E.coli* WK6 (su-) cells. Pre-cultures were grown overnight in LB medium containing 100 µg/ml ampicillin, 2% glucose and 1 mM MgCl_2_. TB medium (2.4 % yeast extract, 2 % tryptone, 0.4 % glycerol, 17 mM KH_2_PO_4_, 72 mM K_2_HPO_4_), supplemented with 100 µg/ml ampicillin, 0.1% glucose and 2 mM MgCl_2_ was inoculated with a 1:50 (v/v) dilution of the pre-culture and cells were grown at 37 °C with 190 rpm. Protein expression was induced with 1 mM IPTG at an OD_600nm_ between 0.7-1.2, before lowering the temperature to 25 °C for 18 hours of expression. Cells were harvested by centrifugation at 4 °C and 4000 x g for 20 min. For lysis, by osmotic shock, a pellet of a 1 l culture (with OD600nm = 25) was resuspended with 10 ml TES buffer (0.2 M Tris pH 8, 0.5 mM EDTA, 0.5 M sucrose) for 2 hours on a rotating wheel. Next, 30 ml TES/4 buffer (TES buffer, four times diluted in H_2_O) was added and left on a rotating wheel for 1 h.

The resuspended cell lysate was centrifuged for 30 min at 8000 x g and the supernatant was kept. Approximately 1 ml of Ni-NTA agarose (Qiagen) was utilised for purification of the lysate resulting from 1 l culture. Pre-equilibrated Ni-NTA agarose beads, in 50 mM sodium phosphate, 1 M NaCl, pH 7, were added to the supernatant and left to incubate on a rotating wheel for 1 h at room temperature. Following incubation the beads were washed with 20 ml 50 mM sodium phosphate, 1 M NaCl, 10 mM imidazole, pH 7 and protein was eluted with 2.5 ml 50 mM sodium phosphate, 0.15 M NaCl, 0.3 M imidazole, pH 7. The elution fractions were dialysed (Spectra/Por 3, 3.5 kDa cut-off) for 3 days against 50 mM HEPES, 0.15 M NaCl pH 7.5 and subsequently concentrated (Amicon, 3 kDa cut-off) to concentrations between 150 µM to 500 µM prior to storage at −80 °C.

### Biophysical analysis

#### SEC-MALS

The oligomerisation states of GBP1 at 15 µM in the presence and absence of GTP and nucleotide analogs were estimated using analytical size exclusion chromatography coupled to multi-angle light scattering (SEC-MALS). Purified protein samples were resolved on a Superdex200 Increase 10/300 GL column (GE Healthcare) connected to a high-performance liquid chromatography (HPLC) unit (1260 Infinity II, Agilent) running in series with an online UV detector (1260 Infinity II VWD, Agilent), an 8-angle static light scattering detector (DAWN HELEOS 8+; Wyatt Technology), and a refractometer (Optilab T-rEX; Wyatt Technology).

For SEC-MALS measurements, proteins were diluted to a final concentration of 15 µM in SEC buffer (50 mM HEPES pH 7.4, 150 mM NaCl, 0.5 mM TCEP or DTT) with or without the GTP transition state mimic or with 1 mM of GTP, GDP, GMP and 0.5 mM of GppCp, GTPyS or GppNHp and incubated for 5 - 10 min at RT prior to injection. On the basis of the measured Rayleigh scattering at different angles and the established differential refractive index increment of value of 0.185 ml*g*^−^*^1^ for proteins in solution with respect to the change in protein concentration (dn/dc), weight-averaged molar masses for each species were calculated using ASTRA software (Wyatt Technology; v.7.3.1).

### Mass photometry

GBP1-WT and GBP1 variants were purified as described before. The data was collected on a Refeyn OneMP instrument using the AcquireMP software (version 2.3 and 2.4). Silicon gaskets (Culture Well Reusable gaskets, Grace Biolabs) were adhered to clean cover slips (High Precision cover slips, No. 1.5, 24 × 50 mm, Marienfeld). For measurements, samples were diluted in 50 mM HEPES pH 7.5, 150 mM NaCl to final concentrations between 12.5 nM to 75 nM. Data was acquired and analysed using DiscoverMP (version 2.3 and 2.4), using the smallest acquisition window and default settings.

### GTPase activity assay

To determine the GTPase activity of GBP1 the GTPase-Glo™ Assay (Promega) was utilised (50), using the protocol for intrinsic GTPase activity. Briefly, 5 µl of 5 µM GBP1 (WT or variants) in running buffer was added per well to a 384 well plate. 5 µl of 2 x GTP solution containing 10 µM GTP and 1 mM DTT was added to the same well. The reaction was incubated at RT for 60 min. 10 µl of reconstituted GTPase-Glo reagent was added to the reaction and incubated for 30 min at RT while shaking. Finally, 20 µl of detection reagent was added and after another incubation step of 10 min the luminescence was measured using a micro plate reader (Synergy*^T^ ^M^*H1, BioTek). BSA was used as a negative control and measurements were performed in triplicates.

### Liposome preparation

#### SUV preparation

1 mg of brain polar lipid extract (BPLE, Avanti Polar Lipids) and 1,2-dioleoyl-sn-glycero-3-phosphocholine (DOPC, Avanti Polar Lipids), purchased as chloroform solutions were each dried under a gentle N2 stream. The resulting lipid film was further dried in a desiccator connected to a vacuum pump for 1 h. To hydrate the lipid film, 1 ml of 50 mM HEPES (pH 7.5), 150 mM NaCl was used. Small unilamellar vesicles (SUVs) were prepared with an Avanti Mini Extruder (Avanti Polar Lipids) with hydrophilic polycarbonate membranes with a pore size of 0.1 µm. The solution of swollen lipid was filled into one of the syringes and monodisperse emulsions of SUVs were produced by passing this mixture through the membrane at least 11 times. The SUVs were stored 4 °C until further use.

#### GUV preparation

Per experimental condition, 30 µl of 10 mg/ml BPLE (Avanti Polar Lipids), was added to 10 µl of 0.1 mg/mL Texas Red 1,2-Dihexadecanoyl-sn-Glycero-3-Phosphoethanolamine, Triethylammonium (Texas red DHPE, Invitrogen) and 1 % (v/v) DSPE-PEG (2000)-biotin (Sigma Aldrich). 30 µl of this solution was carefully aspirated and spread onto a Polyvinyl alcohol (PVA) coated glass cover slide (5 % PVA was prepared in water, dried on a 22x22 mm cover slide for 30 min at 50 °C), prior to an additional 30 min in a desiccator connected to a vacuum pump. To the dried lipid film 250 µl of inside buffer (50 mM HEPES (pH 7.5), 150 mM NaCl, 50 mM sucrose) were added and lipids were allowed to swell in the dark for 15 min with gentle shaking. The giant unilamellar vesicles (GUVs) were collected and freshly used.

### Confocal fluorescence microscopy

#### Preparation of the imaging chamber

Glass coverslips (22x40 mm) were attached, with UV resin, to a homemade pre-drilled piece of plexiglass, to form the imaging chambers. The chambers were flushed with 2 mg/mL BSA-biotin, containing 3 moles of biotin per mole of BSA (BioVision). After removal of Biotin, the chambers were washed with buffer (50 mM HEPES (pH 7.5), 150 mM NaCl) and incubated further 5 minutes with 1 mg/mL Avidin (Thermo Fisher) prior to addition of the GUVs for imaging.

#### Maleimide labelling of GBP1_F_ -Q577C

Alexa Fluor 647 (Thermo Fisher) dissolved in DMSO to a final concentration of 10 mM was added dropwise to the protein until a 20x molar excess was achieved. Prior to addition of the fluorophore, the protein was reduced for 5 min with 0.5 mM TCEP. After addition, the sample was incubated 2 h at room temperature. Separation of the labelled protein from excess dye was performed according to the manufacturer using a desalting column (5 ml, HiTrap Desalting, Cytiva) in 50 mM HEPES pH 7.4, 150 mM NaCl and 0.5 mM.

#### GBP1_F_ -GDP·AlF_3_

The GTP transition state mimic was prepared as described before. 20 µl of Texas red DHPE labelled GUVs were mixed with 5 µl protein solution consisting of 18.5 µM GBP1*_F_* -GDP*·*AlF_3_ and 1.5 µM of GBP1*_F_* - Q577C-GDP*·*AlF_3_, labelled with Alexa 647-C2-maleimide, resulting in a final protein concentration of 4 µM. The mixture was incubated at 30 °C for 30 minutes prior to imaging.

#### GTP-activated GBP1

20 µl of GUVs were mixed with 5 µl protein solution consisting of 18.5 µM GBP1*_F_* and 1.5 µM of GBP1*_F_* -Q577C, labelled with Alexa 647-maleimide. 5 µl of 10 mM GTP was added to the well directly prior to imaging.

#### Confocal microscopy

Imaging was performed on a Nikon A1R confocal microscope using a Nikon SR Apo TIRF 100x oil/1.49 NA objective. The excitation wavelength of the lasers was 561 nm (for Texas Red DHPE) and 640 nm (for GBP1-AF647). Images were processed with Fiji software.

### Dual trap bead-supported membrane transfer assay

#### Bead-supported bilayer preparation

Lipid bilayer coated silica beads were prepared by mixing lipids in chloroform in the desired molar ratios: 84.69 mol% DOPC (850375, Avanti), 15 mol% DOPS (840035, Avanti), 0.15 mol% 18:1 Liss Rhodamine PE (810150, Avanti), 0.16 mol% Biotin lipids DSPE-PEG(2000) Biotin (880129, Avanti).

Lipids were dried to a thin film on the walls of a flask. After removal of residual chloroform, the flask was wrapped in aluminum foil and placed in a desiccator overnight. Lipids were resuspended in 1 mL deionised H_2_O (dH_2_O) to a final lipid concentration of 1 mg/ml, then resuspended for 30 min in a 37°C water bath and subsequently subjected to three freeze-thaw cycles. The lipid solution was extruded 21x using an Avestin-LF-1 extruder with a 100 nm membrane. 10 µL of the lipid solution was added to 89.5 µL H_2_O containing NaCl and 0.5 µL of a 5% solids solution of 2 µm silica microparticles (Sigma Aldrich) to a final concentration of 0.1 mM lipids and 1 mM NaCl, followed by incubation on a lab rotator at room temperature for 45 min. Finally, 30 µL of the solution was diluted into 270 µL GBP buffer (50 mM HEPES pH 7.5, 150 mM NaCl, 5 mM MgCl_2_ and 0.5 mM DTT). For the catching bead, 10 µL of a 200 mM NaCl solution was mixed with 1 µL NTV-DNA 3.5 kDa and 2 µL antiDIG-beads (QDIGP-20-2 ProSciTech), followed by incubation for 30 min on a lab rotator at 4°C and finally diluted in 290 µL GBP buffer.

#### Optical trapping and confocal microscopy

Optical trapping experiments were performed in a LUMICKS C-Trap. One bead containing bead-supported bilayer and one uncoated bead were successively trapped in one of the LUMICKS C-Trap lasers. The bead pairs were then moved far down-stream close to the upper wall of the flow chamber to ensure straight, laminar flow upon switching of the running solution. The bead pairs were flushed at least 30 seconds with GBP buffer + 1 mM GTP at 0.1 bar before dispensing sample containing GBP buffer + 100 µM GBP1 + 1 mM GTP into the flow chamber. A 532 nm laser operated at 8 mW was used to excite 18:1 Liss Rhodamine PE and confocal fluorescence images were acquired in a 14.15x3.35 µm window at 50 nm pixel size to measure Liss Rhodamine PE lipid fluoroescence. The red channel (638 nm) was used to visualise the beads and as internal control for potential dirt particles in the flow channel. Prior to adding GBP1*F*, the beads were flushed for at least 30 seconds with GBP1 buffer supplemented with 1 mM GTP. The time of solute arrival at the first bead was calibrated with a fluorescent dye in separate experiments and estimated to 9 s. To determine fluorescence intensity time traces, the z-axis profile of the inter-bead space and the catch bead was selected in ImageJ and the total relative fluorescence of each frame was processed in Origin. For baseline correction the average fluorescence across 30 seconds prior to addition of GBP1*F* was subtracted from each data set, resulting in a baseline of 0 A.U. All curves were averaged with the “Average multiple curves” option in Origin and the resulting average time traces with the 95% confidence interval were plotted using Python’s matplotlib library.

### Liposome co-sedimentation assays

Wild-type GBP1 and GBP1 variants were *in vitro* farnesylated as described above. Experiments were also performed with *in vivo* farnesylated GBP1 for comparison. The GTP transition state mimic was prepared as described before. GBP1-WT or GBP1 variants were diluted to 2 µM and mixed with 1 mg/ml BPLE SUV liposomes in SEC buffer to a final volume of 100 µl. Samples were incubated for 60 min at RT, followed by ultracentrifugation (Beckman Coulter Optima L-90K, Rotor: 42.2 Ti) at 222,654 x g for 20 min at 4°C. The pellet and supernatant fractions were separated as quickly and gently as possible prior to analysis, whereby 15 µl were loaded onto a 4-12 % SurePAGE Bis-Tris gel for separation by SDS-PAGE.

The lanes of interest were identified and the bands automatically detected using the Gel doc Image Lab software. After automatic background subtraction, the sum of intensities of the protein present in the pellet and supernatant fractions was used to determine the relative percentage of GBP1 in each fraction. The pelletation assay was performed five to seven times to compute fractional average intensities and standard deviations.

### Negative staining EM

3.5 µL of protein or lipid solution was applied onto a freshly glow-discharged carbon-coated copper mesh grid (Quantifoil). After 1 min, grids were washed twice with 12 µL buffer (50 mM HEPES pH 7.4, 150 mM NaCl, 0.5 mM DTT) followed by staining with 3.5 µL of 2 % (w/v) uranyl acetate at room temperature. At each step, excess sample, wash solution and stain were blotted with filter paper and finally grids were air dried for 15 min. Grids were imaged on a JEM 1400Plus TEM (JEOL) operated at 120 kV and recorded on a bottom-mounted TVIPS F416 CMOS camera.

To observe micelle formation, 0.5 - 2 mg/ml of GBP1*_F_*- GDP*·*AlF_3_ or 8 mg/ml of GBP1*_F_* with 1 mM of GTP was applied onto a carbon grid. The incubation time for GBP1*_F_*- GDP*·*AlF_3_ was 10 min at RT and 2 min at RT for GTP respectively. To analyse membrane binding of GBP1*_F_*, 1 mg/ml of GBP1*_F_* was added together with all other components to form the GTP-transition state mimic (see above). SUVs were added in a 1:10 dilution (10 mg/ml, d = 100 nm and incubated for 10 min at RT. For filament formation to occur, samples needed to incubate o/n at 4°C, 30 min at 30°C or the concentration needed to increase to 2 mg/ml following an incubation step of 10 min at RT.

### Single particle imaging

#### GBP1-GDP·AlF_3_ dataset

A total of 3.0 µL of 0.7 mg/ml GBP1-GDP*·*AlF_3_ was applied to glow-discharged Quantifoil grids (QF-1.2/1.3, 300-mesh holey carbon on copper) on a Leica GP2 vitrification robot at 99 % humidity and a temperature of 22 °C. The sample was blotted for 4 s from the carbon side of the grid and immediately flash-cooled in liquid ethane. Micrographs were acquired on a FEI Titan Krios (Thermo Fisher Scientific) operated at 300 kV. Images were recorded on a K2 Summit direct electron detector (Gatan) with a pixel size of 1.09 Å. Image acquisition was performed with EPU Software (Thermo Fisher Scientific), and micrographs were collected at an underfocus varying between −3.5 µm and −0.5 µm. We collected a total of 48 frames accumulating to a total exposure of 60 e*^−^*/Å^2^. In total, 1,193 micrographs were acquired. Data acquisiton parameters are summarised in Extended Data Table S2.

#### GBP1-GDP·AlF_3_-Nb74 dataset

GBP1 was incubated in a 1:1 molar ratio with Nb74 for 70 min at RT, before adding GDP*·*AlF_3_ and incubating additionally for 10 min at RT. A total of 3.0 µL of 0.7 mg/ml GBP1-GDP*·*AlF_3_ bound to Nb74 was applied to glow-discharged Quantifoil grids (QF-1.2/1.3, 300-mesh holey carbon on copper) on a GP2 vitrification robot at 99 % humidity and 22 °C. The sample was blotted for 4 s from the carbon side of the grid and immediately flash-cooled in liquid ethane. Micrographs were acquired on a FEI Titan Krios (Thermo Fisher Scientific) operated at 300 kV. Images were recorded on a K3 Summit direct electron detector (Gatan) at a magnification of 105kx, corresponding to a pixel size of 0.834 Å at the specimen level. Image acquisition was performed with EPU 2.8.1 Software (Thermo Fisher Scientific), and micrographs were collected at an underfocus varying between −2.2 µm and −0.6 µm. We collected a total of 50 frames accumulating to a total electron exposure of 60 e*^−^*/Å^2^. In total, 5,214 micrographs were acquired.

#### GBP1_F_ -GDP·AlF_3_

A total of 3.0 µL of 1 mg/ml GBP1*_F_* - GDP*·*AlF_3_ was applied to glow-discharged Quantifoil grids (QF-1.2/1.3, 300-mesh holey carbon on copper) on a Leica GP2 vitrification robot at 97 % humidity and 20 °C. The sample was blotted for 4 s from the carbon side of the grid and immediately flash-cooled in liquid ethane. Grids were imaged on a JEM 3200FSC TEM (JEOL) operated at 300 kV. Images were recorded on a K2 Summit direct electron detector (Gatan). Two different datasets were acquired one at a magnification of 30kx, corresponding to a pixel size of 1.22 Å at the specimen level and the other one at a magnification of 15kx, corresponding to a pixel size of 2.449 Å. Image acquisition was performed with SerialEM (51), and micrographs were collected at an underfocus varying between −3 µm and −1 µm. We collected a total of 60 frames accumulating to a total electron exposure of 48.37 e*^−^*/Å^2^ (for the dataset at 30kx) or to a total electron exposure of 12.92 e*^−^*/Å^2^ (for the dataset at 15kx). In total, 395 (dataset at 30kx) or 606 (dataset at 15kx) micrographs were acquired.

#### GBP1_F_ -GDP·AlF_3_ with LPS from E.coli O111:B4 (LPS-EB), Salmonella Minnesota R595 (LPS-SM) or from Salmonella Typhimurium (LPS-ST)

1 mg/ml GBP1*_F_* -GDP*·*AlF_3_ was mixed with 0.22 mg/ml LPS-EB (InvivoGen), LPS-SM (InvivoGen) or LPS-ST (Enzo) and incubated for 30 min at 30 °C. 3.0 µL of the mixture were applied on to glow-discharged Quantifoil grids (QF-1.2/1.3, 200-mesh holey carbon on copper) on a Leica GP2 vitrification robot at 98 % humidity and 22 °C. The sample was blotted for 4 s from the carbon side of the grid and immediately flash-cooled in liquid ethane. Grids were imaged on a JEM 3200FSC TEM (JEOL) operated at 300 kV. Images were recorded on a K2 Summit direct electron detector (Gatan) using automated image acquisition in SerialEM (51). Data colelction statistics for each dataset are summarised in Extended Data Table S4.

### Single-particle image processing

#### GBP1-GDP·AlF_3_-Nb74

The GBP1-GDP*·*AlF_3_-Nb74 dataset was processed using cryoSPARC v3.3.2 (52). The in-built patch-motion correction (53) routine in cryoSPARC was used to correct for stage drift and beam-induced specimen movement over the acquired frames. 5,208 micrographs were selected for further processing and patched contrast transfer function (CTF) determination (54) was performed in cryoSPARC. Using a blob-based particle picker, 2,171,521 particles were extracted and cleaned via multiple rounds of 2D classification, each consisting of 50 - 100 classes. Classes only containing the LG domain of the protein were actively sorted out as they did not yield a full 3D reconstruction [Extended Data Fig. S21]. Selected 2D classes comprising 119,071 particles were used to train a Topaz model (55), which was then used to extract a total of 6,539,167 particles. Following particle extraction, four iterative rounds of 2D classification were performed and 2D class averages were selected displaying secondary structure features. 500,186 particles were used to perform *ab initio* reconstruction to generate five different models. Three classes were selected for heterogeneous refinements without imposing symmetry or imposing C2 symmetry. A single class with 187,161 particles was selected and used for non-uniform refinement (56) either without imposing symmetry or with imposed C2 symmetry [Extended Data Fig. S4]. Per-particle defocus and global CTF refinement improved the resolution to 3.7 Å. Local resolution was estimated in cryoSPARC (57) and visualised in ChimeraX (58). Map sharpening was done in cryoSPARC by applying the overall B-factor estimated from Guinier plots. For flexible refinement, the final particle stack of the cryo-EM density was clipped to 256 pixel, fourier-cropped to 96 pixel (pixel size 2.2240 A) and used as input for cryoSPARC v4.1.1 3D Flexible Refinement ((59)) with 4 latent dimensions. Morphs of the density along the four dimensions of the latent space were generated to display different modes of flexibility and were displayed in ChimeraX (58).

#### GBP1-GDP·AlF_3_

1,193 movies of GBP1-GDP*·*AlF_3_ were processed in cryoSPARC v3.1 (52). Patch-motion correction and patched CTF estimation were followed by manual particle picking. Those manual picks were used to train a Topaz model (55) from which 240,487 particles were extracted. After multiple rounds of 2D classification, particles assigned to classes displaying secondary structure were used as an input to perform *ab initio* reconstruction to generate 5 different models (67,197 particles). Three classes were used for heterogeneous refinement imposing C2 symmetry. A final non-uniform refinement (56) consisting of 35,715 particles resulted in a 4.9 Å resolution structure that only covered the LG domain of GBP1.

#### GBP1-GDP_F_ ·AlF_3_

Images of GBP1-GDP*_F_ ·*AlF_3_ micelles were processed in cryoSPARC v3.1.0 (52). Patch-motion correction and patched CTF estimation were followed by manual particle picking and 2D classification.

#### GBP1-GDP_F_ ·AlF_3_ with LPS-EB

The dataset was processed using cryoSPARC v3.3.2 (52). After patch-motion correction and patched CTF estimation, the particle segments were generated from traced filaments using the cryoSPARC filament tracer. After multiple rounds of 2D classification, classes displaying clear molecular features were used as templates for the cryoSPARC template picker. Extracted particles were again subjected to multiple rounds of 2D classification.

### Atomic model building

Atomic models of the GTPase Domain of human GBP1-GDP*·*AlF_3_ (PDB ID 2b92) (37) and the C-terminal stalk (aa 320 - 483) of GBP1 (PDB ID 1dg3) (35) were rigid body-fitted into the cryo-EM density. Manual model building was performed in Coot 0.9.5 (60) followed by real-space refinement against one of the half maps in Phenix 1.13 (61). The second half map was used as a test map for assessment of overfitting. Ligand geometry and restraints for GDP*·*AlF_3_ were generated using the electronic Ligand Builder and Optimisation Workbench (eLBOW) (62) implemented in Phenix. Secondary structure restraints were used throughout the refinement. A locally sharpened and filtered map was generated using the hybrid version of LocScale (39), which integrates reference based sharpening for modelled regions (63) with generalised scattering properties of biological macromolecules for unmodelled regions approximated by pseudo-atoms (40). The atomic displacement factors of the combined model were refined using 10 Refmac (64) iterations as implemented in servalcat (65) with the keywords ’refi bonly’. Prior to refinement, all the atomic displacement factors were set to 40 Å^2^.

### Tomography

#### GBP1_F_ -GDP·AlF_3_ with liposomes

For the dataset of GBP1*_F_* -GDP*·*AlF_3_ together with liposomes, a total of 3.5 µL of 1 mg/ml GBP1*_F_* -GDP*·*AlF_3_ containing freshly extruded liposomes (1 mg/ml of BPLE, Avanti, d = 100 nm) and 10 nm gold fiducials (1:5 (v/v)) was applied to glow-discharged Quantifoil grids (QF-1.2/1.3, 200-mesh holey carbon on copper, Quantifoil) on a Leica GP2 vitrification robot (Leica) at 98 % humidity and 20 °C. The sample was blotted for 4 s from the carbon side of the grid and immediately flash-cooled in liquid ethane. Grids were imaged on a JEM 3200FSC TEM (JEOL) operated at 300 kV. Images were recorded on a K2 Summit direct electron detector (Gatan) at a magnification of 12kx, corresponding to a pixel size of 3.075 Å at the specimen level. Image acquisition was performed with SerialEM (51), and micrographs were collected at a nominal defocus of −5 µm or −4 µm. Bidirectional tilt series were acquired from 0° to -60° and from 0° to 60° with a 2° increment. We collected tilt series of 61 micrographs each consisting of 10 frames and a total electron exposure of 93.94 e*^−^*/Å^2^ for tomogram-33 and tomogram-39 (1.54 e*^−^*/Å^2^ per tilt increment). For tomogram-50, we collected a tilt series of 61 micrographs consisting of 20 frames and a total electron exposure of 100.04 e*^−^*/Å^2^ (1.54 e*^−^*/Å^2^ per tilt increment). Micrographs were motion-corrected with MotionCor2 (53) and dose-weighted according to their accumulated electron exposure (66). CTF correction was performed using ctfphaseflip from the IMOD package (67) and the tilt series was aligned using patch tracking and reconstructed using weighted back-projection as implemented in Etomo from the IMOD package. Segmentation of lipid membranes and protein coat was performed with tomoseg as part of the EMAN2 package (68) on reconstructed tomograms binned by a factor of 2. Segmented tomograms were visualised with ChimeraX (58).

### Bioinformatic analysis

#### Multiple sequence alignment

The sequence of hGBP1 - hGBP7 (UniProt: P32455, P32456, Q9H0R5, Q96PP9, Q96PP8, Q6ZN66, Q8N8V2) were used as input for Clustal Omega (69). The resulting sequence alignment was displayed and consensus sequences computed in MView (70).

#### Sequence conservation

The sequence of hGBP1 (UniProt: P32455) was used as input for the ConSurf Server (71) to search the UniRef90 database with the HMMer (72) using one iteration. The resulting sequence alignment was displayed and consensus sequences computed in MView (70). The conservation was mapped onto the atomic model using ChimeraX (58).

## ACKNOWLEDGEMENTS

We thank Roland Kieffer and Jeremie Capulade for help with fluorescence imaging, Mario Avellaneda and Florian Wruck for setting up the initial optical trapping experiments and Wiel Evers for cryo-EM data collection. We acknowledge Instruct-ERIC (PID7267), part of the European Strategy Forum on Research Infrastructures (ESFRI), and the Research Foundation - Flanders (FWO) for their support and use of resources, as well as Alison Lundqvist for technical assistance during nanobody discovery. Cryo-EM data collection benefited from access to the Netherlands Centre for Electron Nanoscopy (NeCEN) with financial support from the Dutch Roadmap Grant NEMI (NWO.GWI.184.034.014). This work was supported by the European Research Council (ERC-StG-852880 to AJ), the Dutch Research Council (NWO.STU.018-2.007 to AJ) and the Kavli Institute of Nanoscience Delft.

## Extended Data Movies

SM1 Movie_SM1.mp4 - 3D flexible refinement of the GBP1:Nb74 complex.

SM2 Movie_SM2.mp4 - Confocal imaging of GBP1*_F_* -dependent membrane fragmentation and lipid transfer.

SM3 Movie_SM3.mp4 - Electron cryotomogram of GBP1 coatomer on BPLE SUVs and close-up of GBP1*_F_* -GDP*·*AlF_3_ micelle. SM4 Movie_SM4.mp4 - Electron cryotomogram of GBP1 coatomer on BPLE SUVs as well as rendering of segmented density of 3D tomographic reconstruction.

---

**Extended Data Fig. S1.**
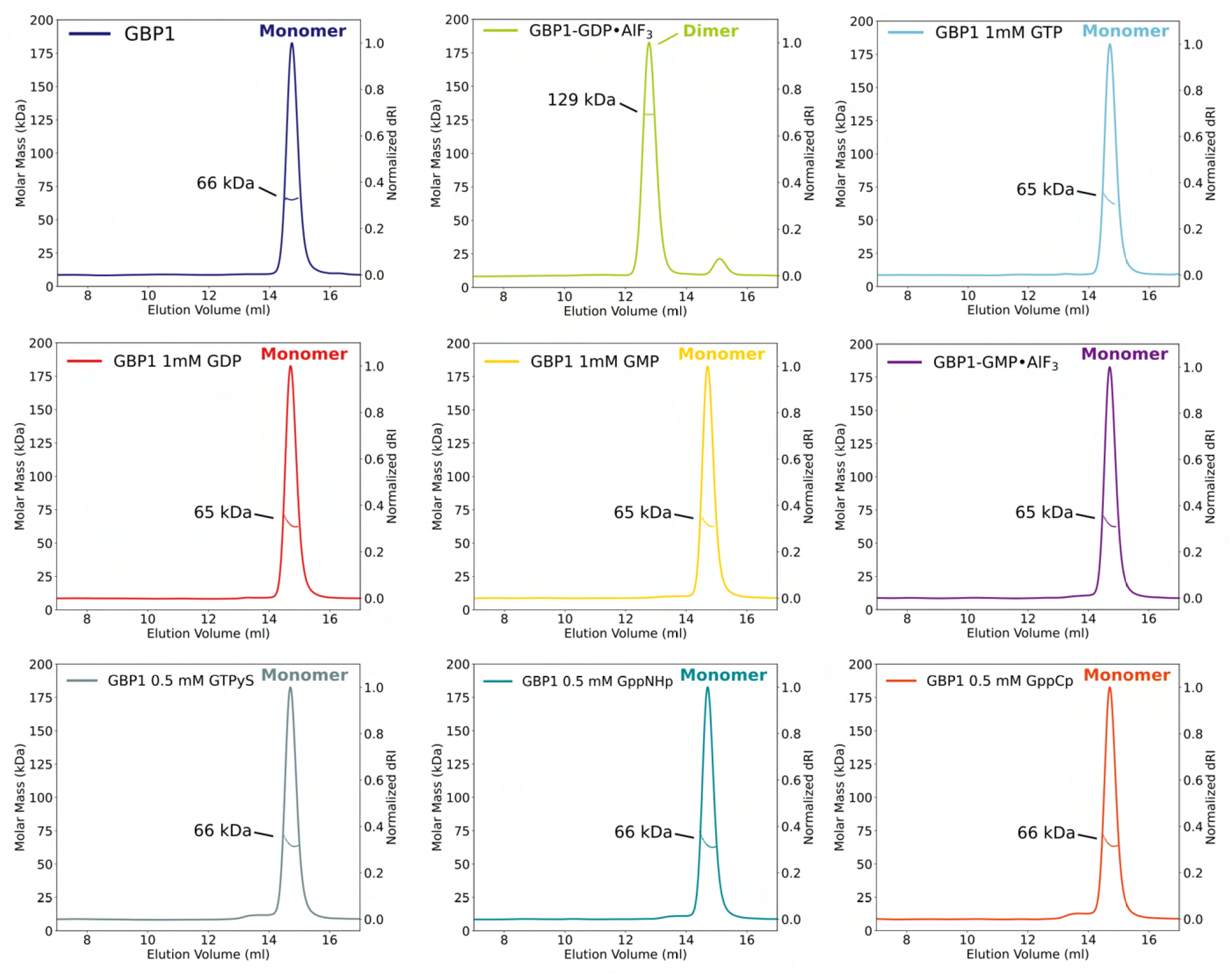
Individual SEC-MALS experiments for GBP1 with different guanine nucleotides. Size exclusion profiles of GBP1 in the presence of GTP, GDP, GMP, GMP*·*AlF3, GTP*γ*S, Guanosine-5’-[(*β*,*γ*)-imido]triphosphate (GppNHp) and Guanosine-5’-[(*β*,*γ*)-methyleno]triphosphate (GppCp) are consistent with a GBP1 monomer, while a GBP1 dimer peak emerges in the presence of GDP*·*AlF_3_ . The experimentally determined molecular weight is plotted across the chromatographic peak and is reported in kDa.

**Extended Data Fig. S2.**
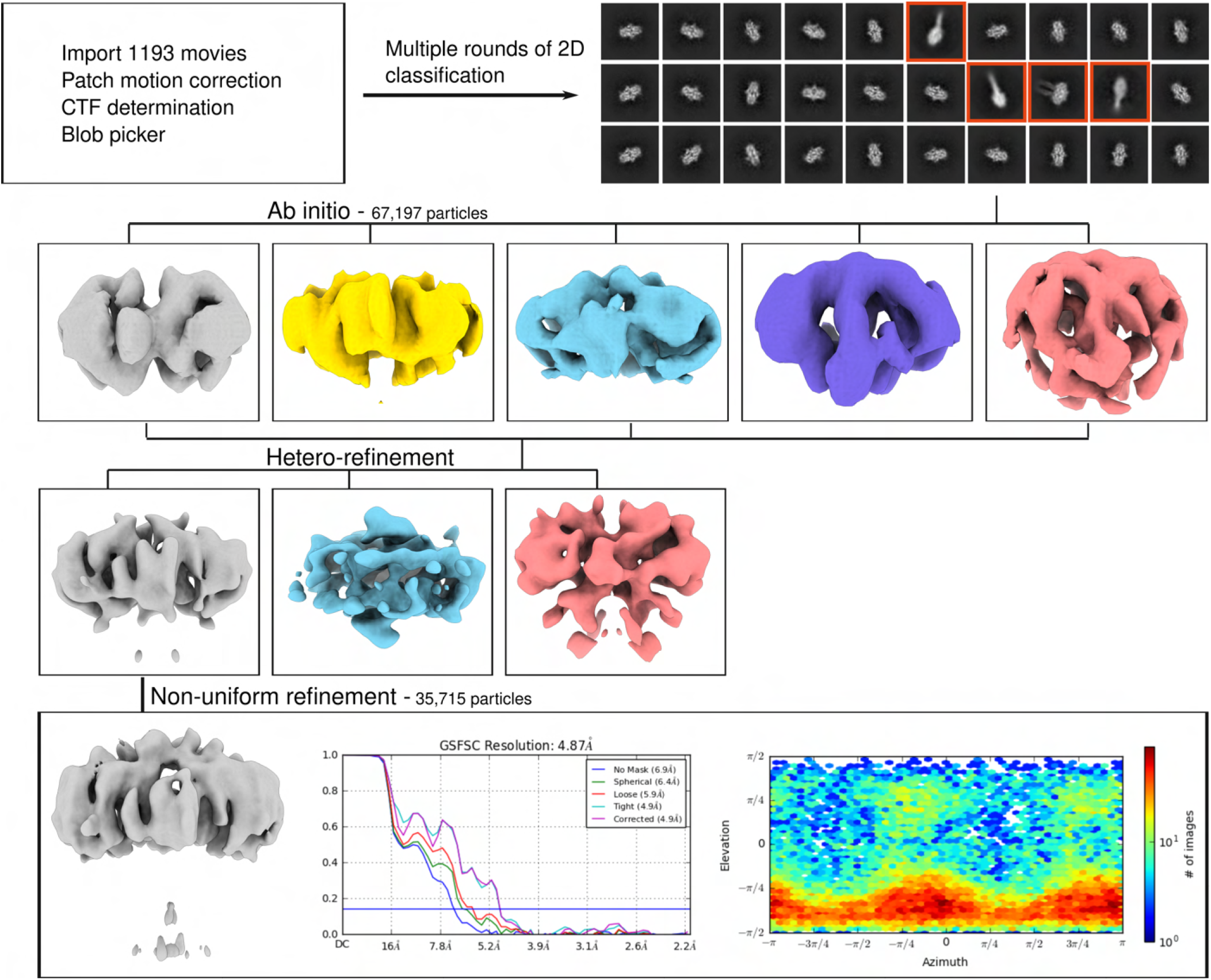
Image processing for GBP1-GDP*·*AlF_3_. Single-particle analysis processing workflow of GBP1-GDP*·*AlF_3_ converged on the LG domain dimer. A majority of 2D classes (92% of all particles) showed a top view representative projection of the LG domain dimer. A subset of 2D classes also comprised the MD domain of both monomers (red boxes).

**Extended Data Fig. S3.**
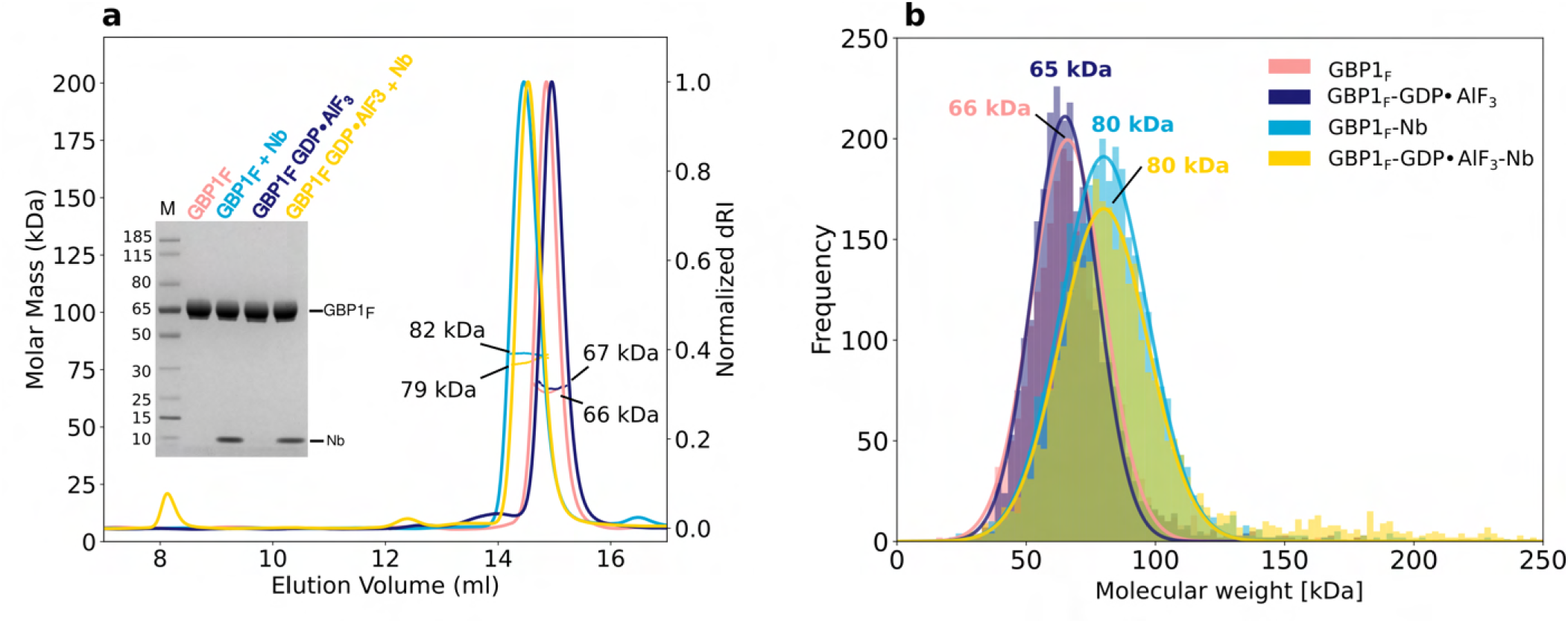
GBP1*_F_* - Nb74 interaction. (a) SEC-MALS experiments showing that farnesylated GBP1 (GBP1*_F_*) appears primarily monomeric in the absence and in the presence of GDP*·*AlF_3_ (MW = 66 kDa and 67 kDa). Nb74 binds GBP1*_F_*with 1:1 stochiometry both in the absence and in the presence of GDP*·*AlF_3_ . For conditions containing GDP*·*AlF_3_, we frequently observed an additional peak close to the void volume of the SEC column corresponding to higher molecular weight species. The inset shows SDS-PAGE analysis of the SEC-MALS input. (b) Mass photometry analysis confirming that GBP1*_F_* is monomeric in the presence of GDP*·*AlF_3_ and the 1:1 stochiometry of Nb74 binding to GBP1*_F_*. Note that rare events corresponding to large GBP1*_F_* assemblies such as those observed by SEC-MALS may not be detected in the chosen field-of-view for the experiments shown

**Extended Data Fig. S4.**
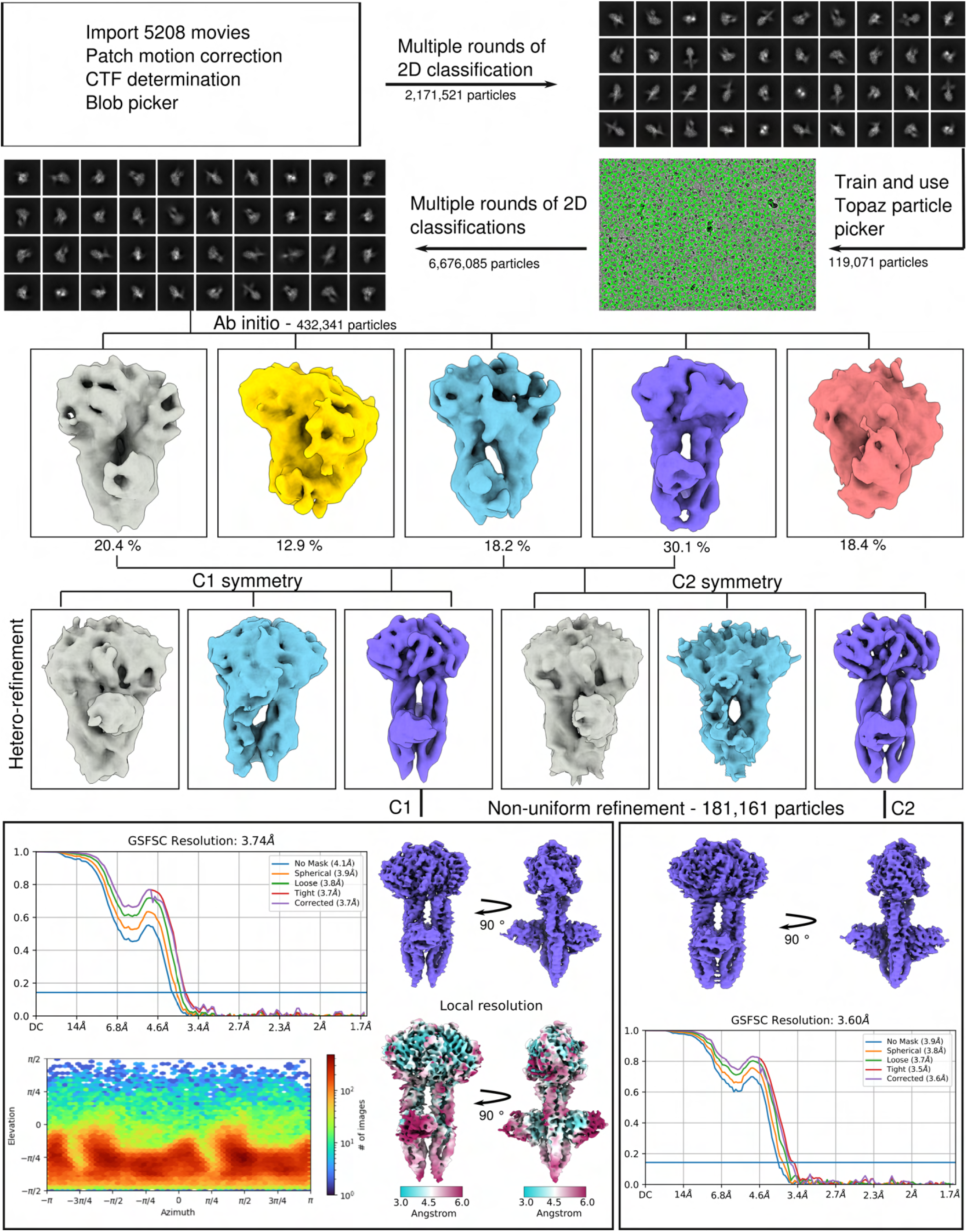
Image processing and structure determination of GBP1-GDP*·*AlF_3_ -Nb74. The image processing workflow is displayed for both the C1 reconstruction and for the reconstruction with C2 symmetry imposed.

**Extended Data Fig. S5.**
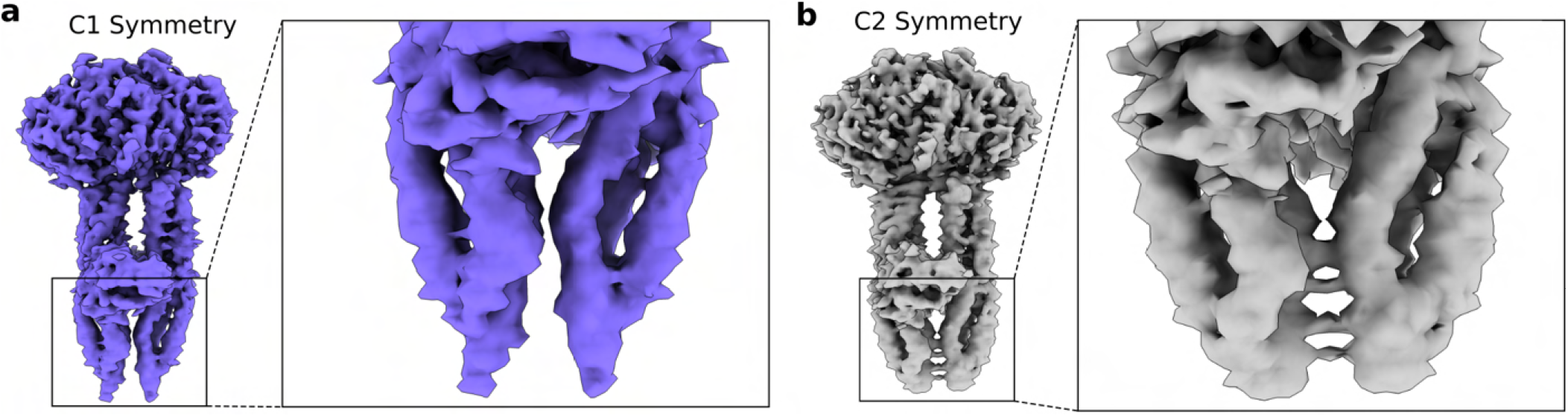
Comparison of MD density in C1 and C2-symmetrised cryo-EM maps of GBP1-GDP*·*AlF_3_-Nb74. (a) Cryo-EM density without imposed symmetry (C1) or (b) with C2 symmetry imposed. Zoom boxes show the C-terminal end of the middle domain (MD). Imposing strict C2 symmetry during reconstruction resulted in symmetrisation artefacts arising at the C-terminal part of the MD domain.

**Extended Data Fig. S6.**
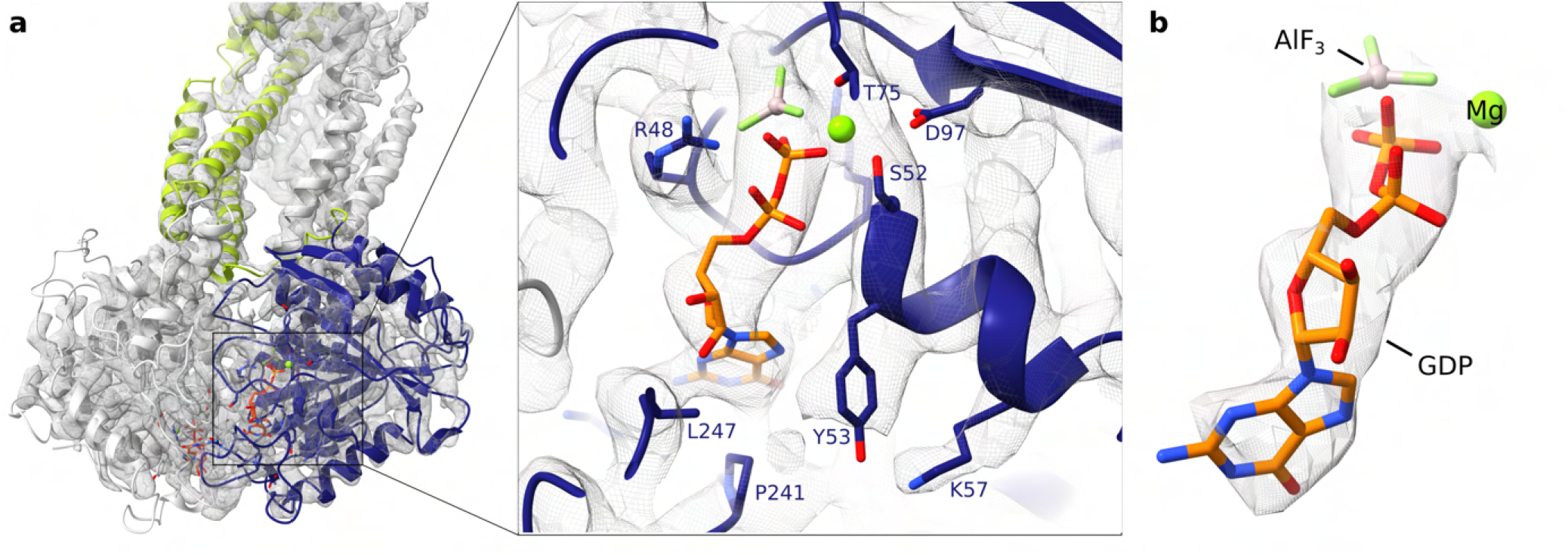
Guanine nucleotide binding site. ((a) Location of one of the two GDP*·*AlF_3_ ligands at the GBP1 dimer interface formed by the LG domains and close-up of the catalytic site with the ligand in stick representation superposed onto the cryo-EM density. The catalyic arginine R48 and residues in close proximity to the ligand are also highlighted in stick representation. (b) GDP*·*AlF_3_ ligand displayed with cropped cryo-EM density.

**Extended Data Fig. S7.**
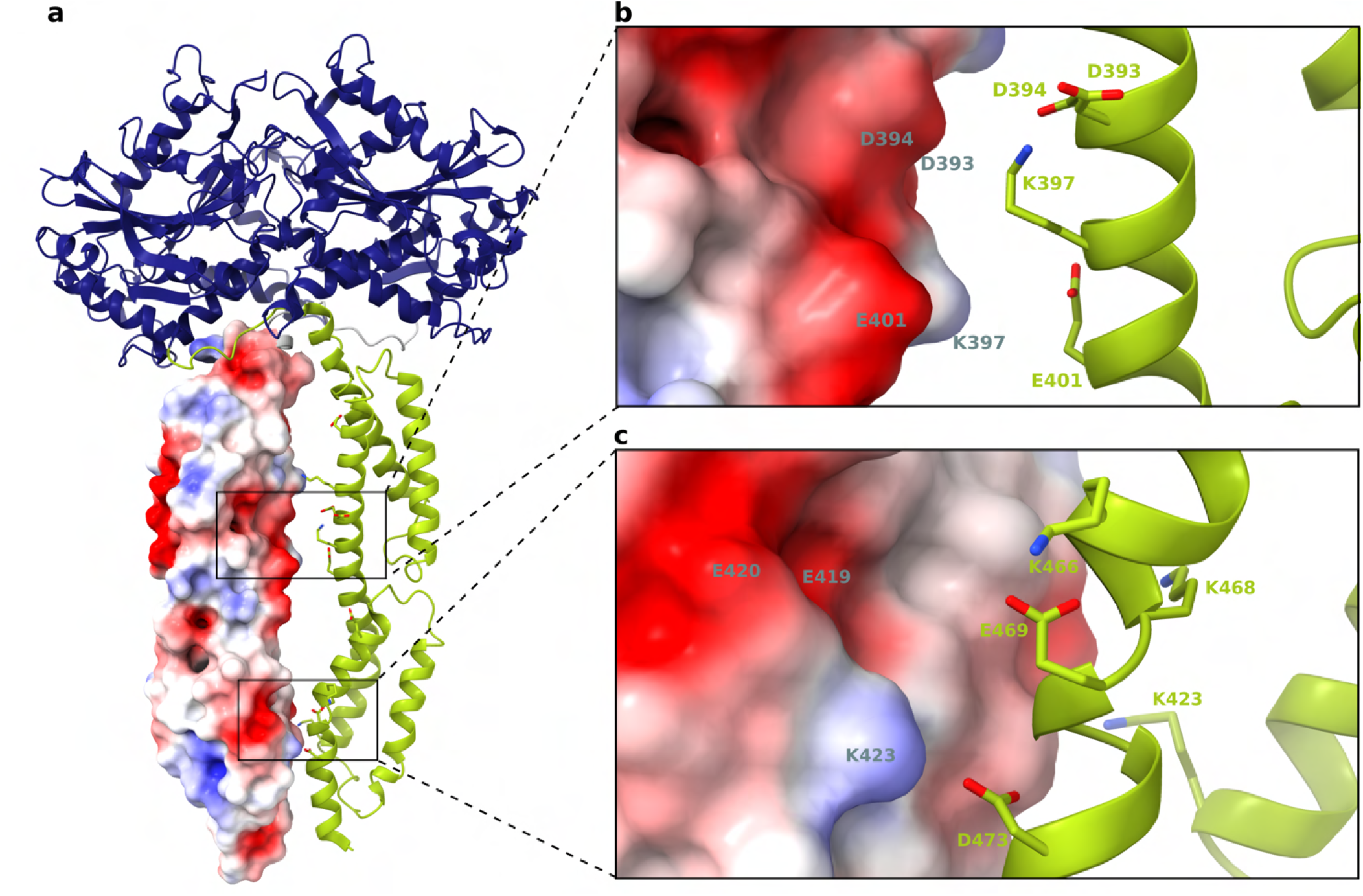
Overview of charged residues in the C-terminal helical domain (CTHD) of the GBP1 dimer. (a) Charged residues at the interface between the CTHDs are displayed as sticks for one of the monomers. The CTHD of the other monomer is displayed in surface representation with its mapped electrostatic potential. Note that the resolution of the EM density map in this region did not allow unambiguous modelling of their side chain conformations and only preferential rotamers are shown without reference to potential interactions. (b) and (c) Close-up of the MD interfaces between two GBP1 monomers in the dimer. Residues with opposing charges can be found on both sides of the interface, potentially stabilising the two parallel middle domains of the GBP1 dimer by electrostatic interactions.

**Extended Data Fig. S8.**
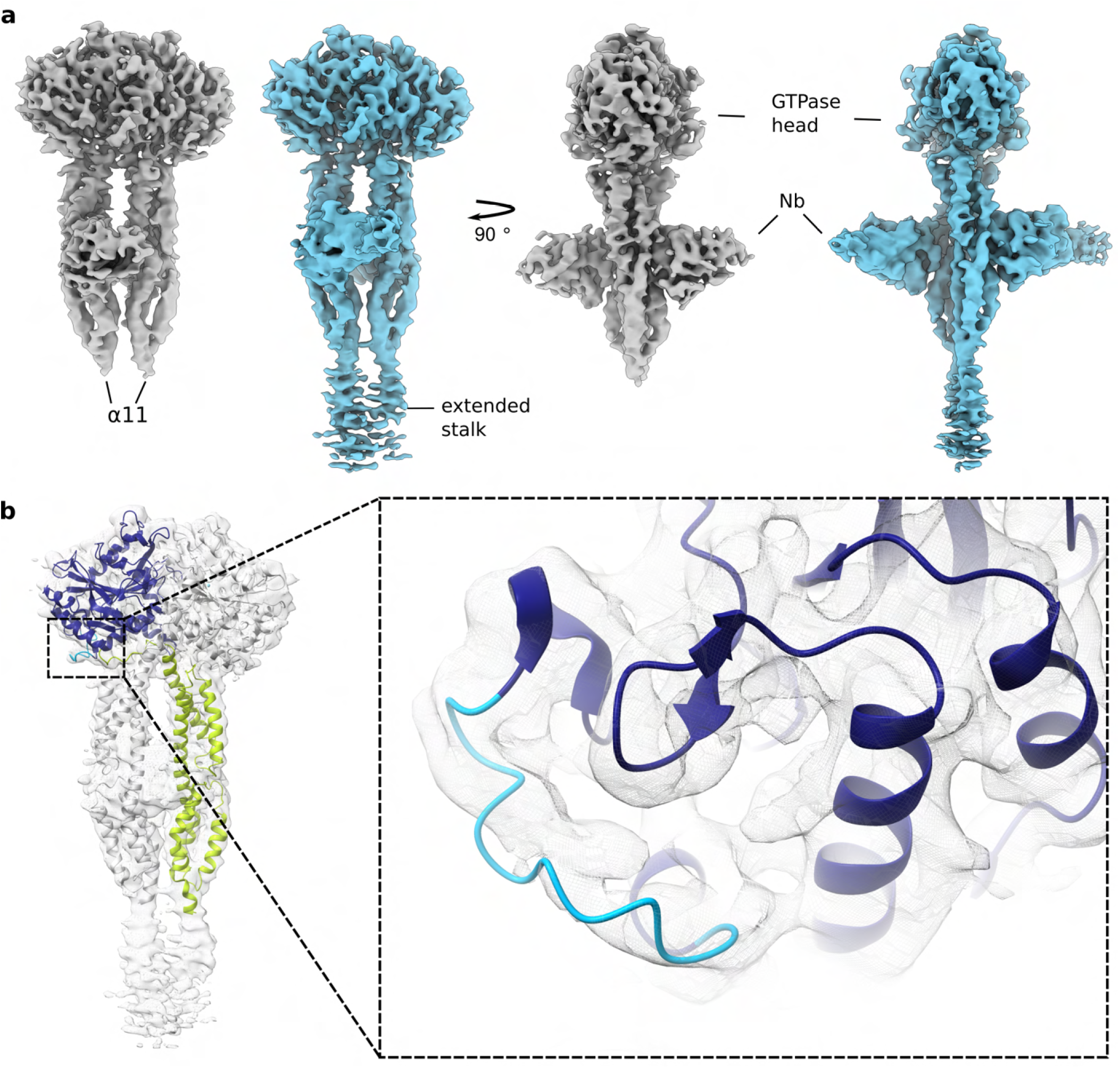
LocScale map of the GBP1–Nb74 dimer. (a) Locally sharpened density maps reveal additional density protruding from helix α11 consistent with a flexible GED. (b) The locally sharpened map also allowed tracing the α_3_-α_3_’ loop (residues 156-167), highlighted in light blue.

**Extended Data Fig. S9.**
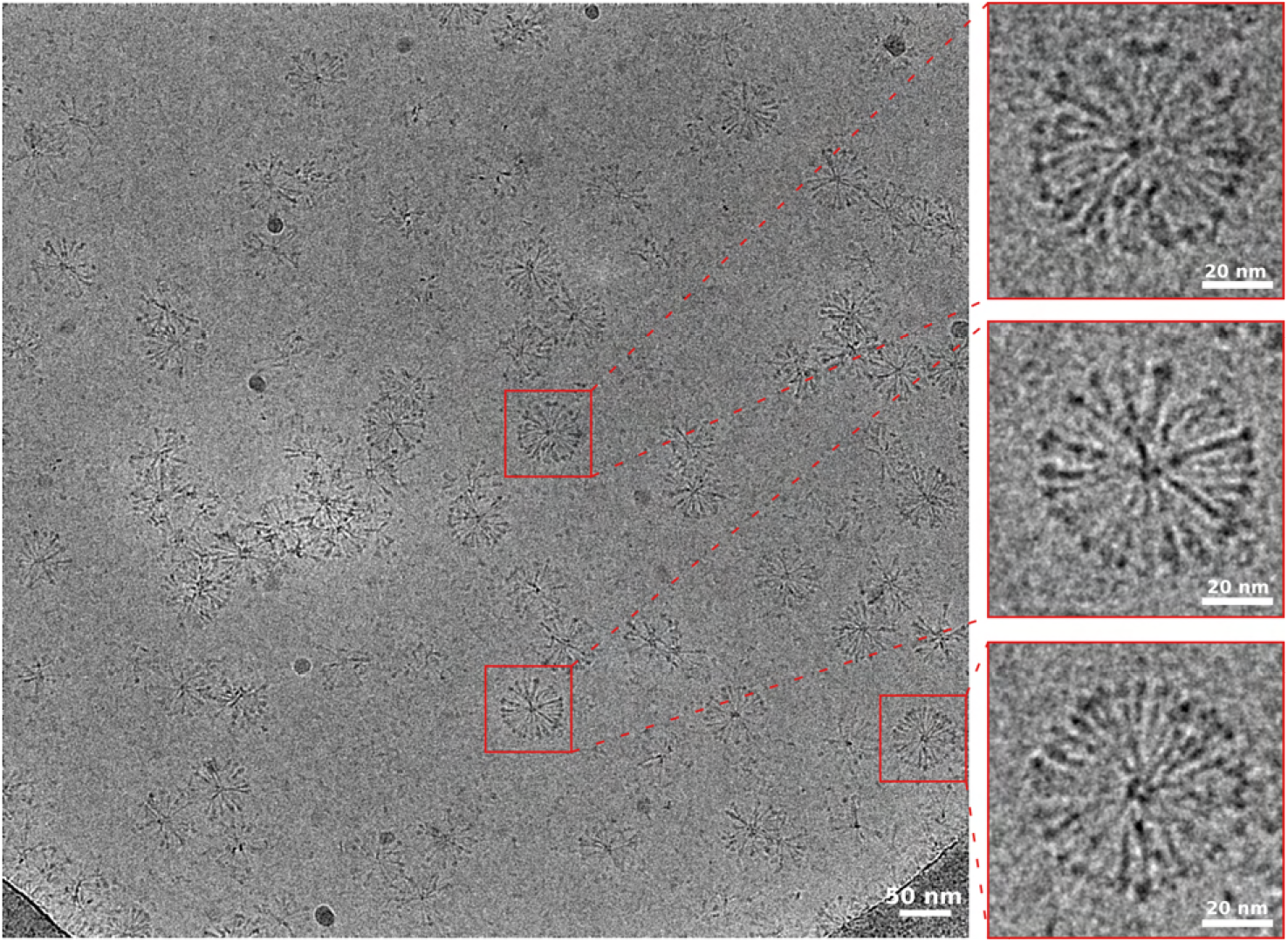
Representative cryo-EM micrograph of GBP1*_F_* in the presence of GDP*·*AlF_3_. In the absence of lipids, farnesylated GBP1*_F_* oligomerises into flower-like assemblies.

**Extended Data Fig. S10.**
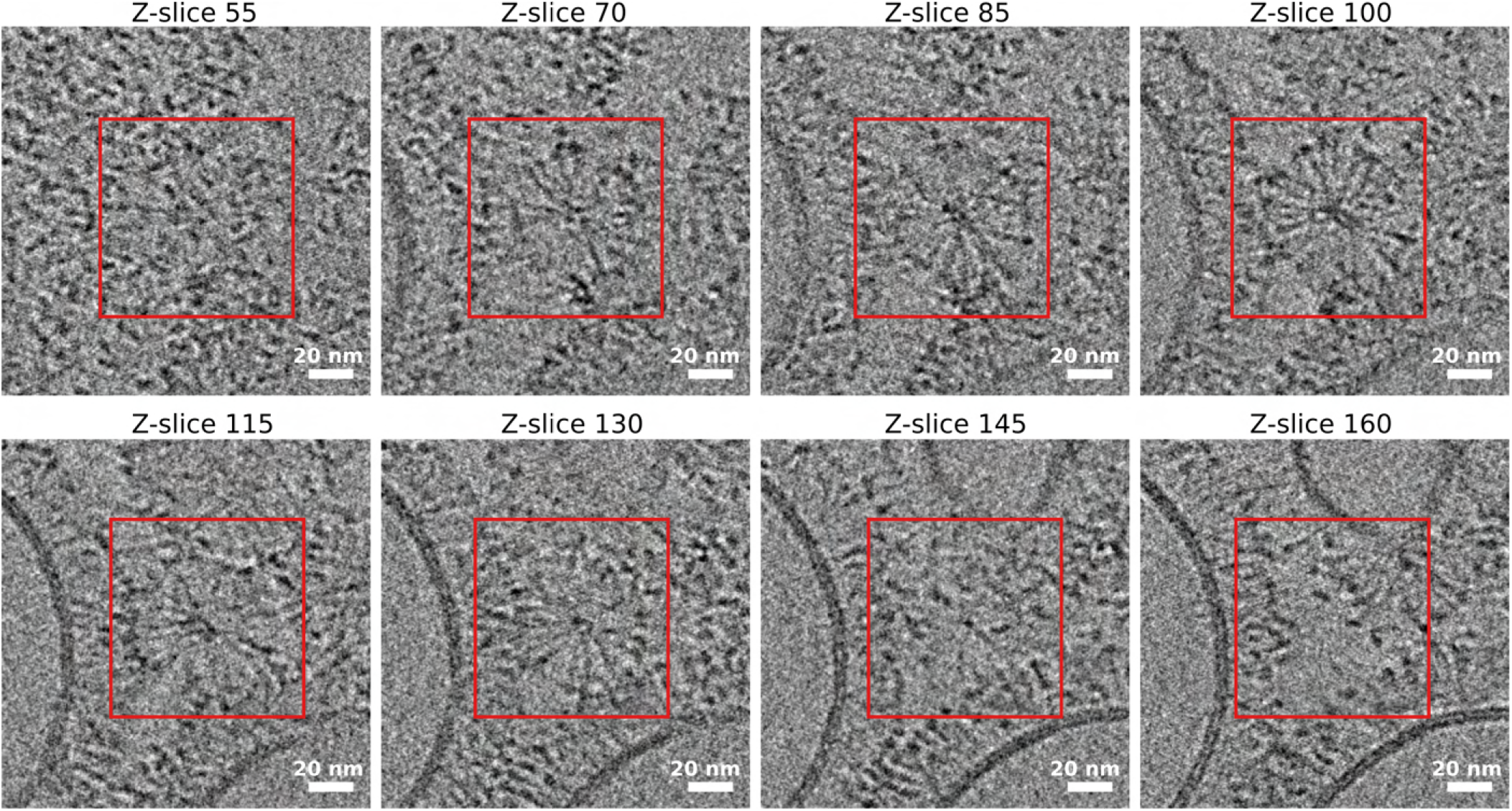
Tomographic z-stack of spherical, micellar GBP1*_F_* assemblies. Slices through the z-stack show the varying diameter of GBP1*_F_*-GDP*·*AlF_3_ assemblies consistent with a spherical geometry. Associated movie: Extended Data Movie SM3.

**Extended Data Fig. S11.**
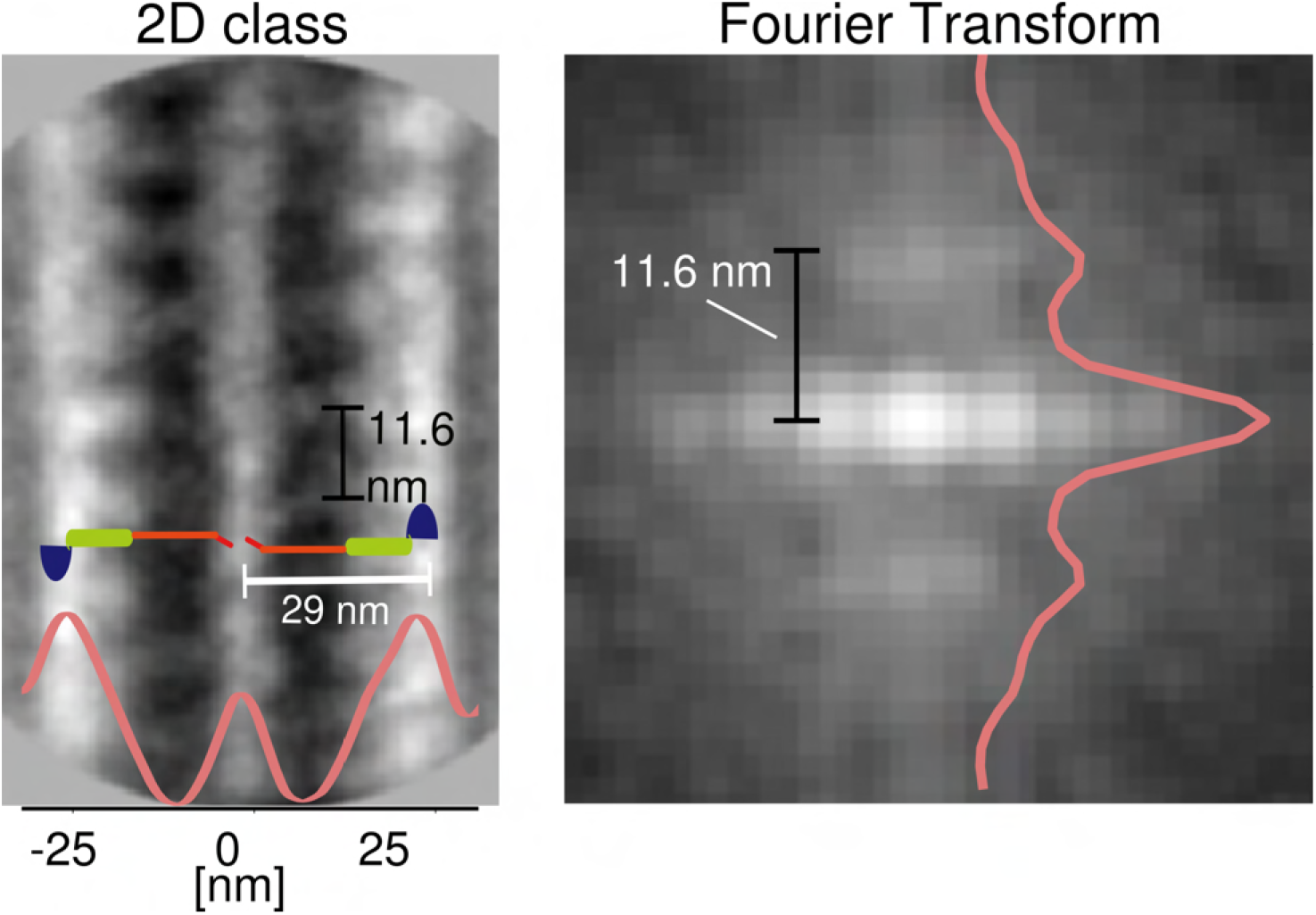
GBP1*_F_*-GDP*·*AlF_3_ tubular protrusion. 2D class average of negatively stained GBP1*_F_*-GDP*·*AlF_3_ tubular protrusion (left panel). The computed power spectrum shows a principal layer line at 0.086 Å^-1^, corresponding to a periodicity of 11.6 nm along the filament axis (right panel).

**Extended Data Fig. S12.**
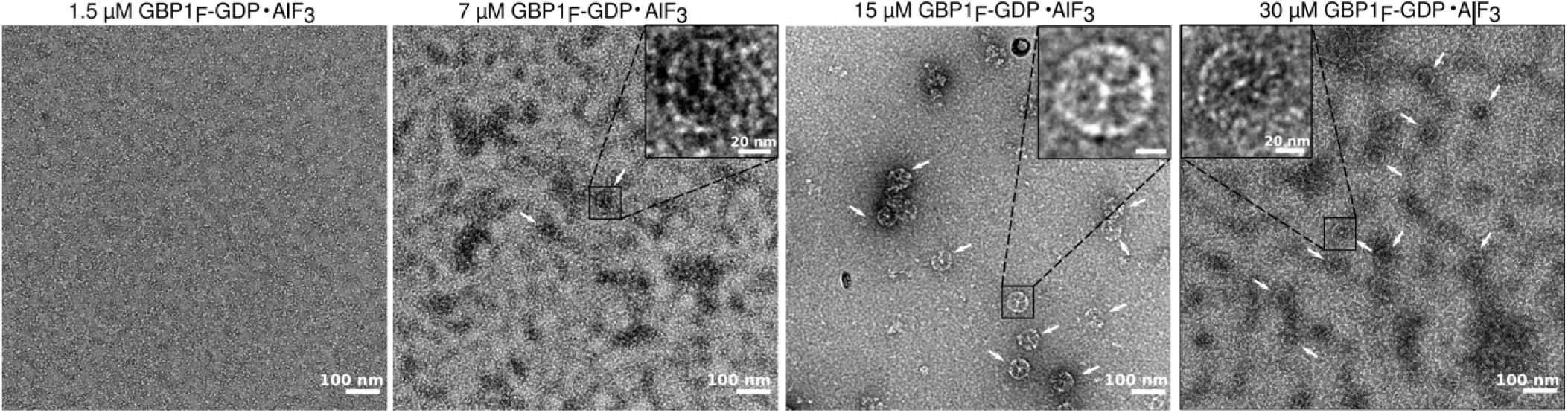
Micelle formation of GBP1*_F_*-GDP*·*AlF_3_ is concentration dependent. Formation of micellar assemblies of GBP1*_F_*-GDP*·*AlF_3_ (white arrows) starts beyond a threshold concentration of around 7 µM.

**Extended Data Fig. S13.**
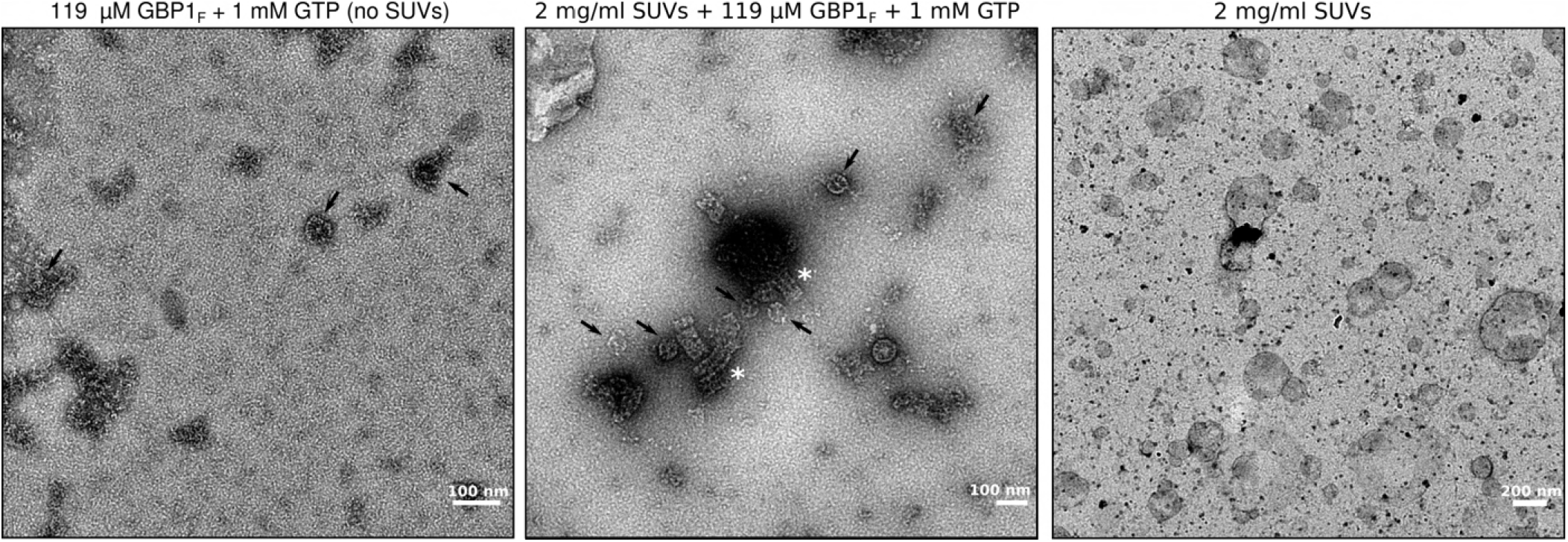
GTP-induced assembly of GBP1*_F_* micelles and membrane fragmentation. Micelle formation of GBP1*_F_* (black arrows) was observed after addition of 1 mM GTP to a highly concentrated GBP1*_F_* solution (119 µM) in the absence of lipids (left panel). Upon addition of 2 mg/ml SUVs, GBP1*_F_* remodelled SUVs into spherical micelles (middle panel; arrows) and short filaments (white asterisks). A sample containing 2 mg/ml of SUVs without GBP1*_F_* is shown for comparison (right panel).

**Extended Data Fig. S14.**
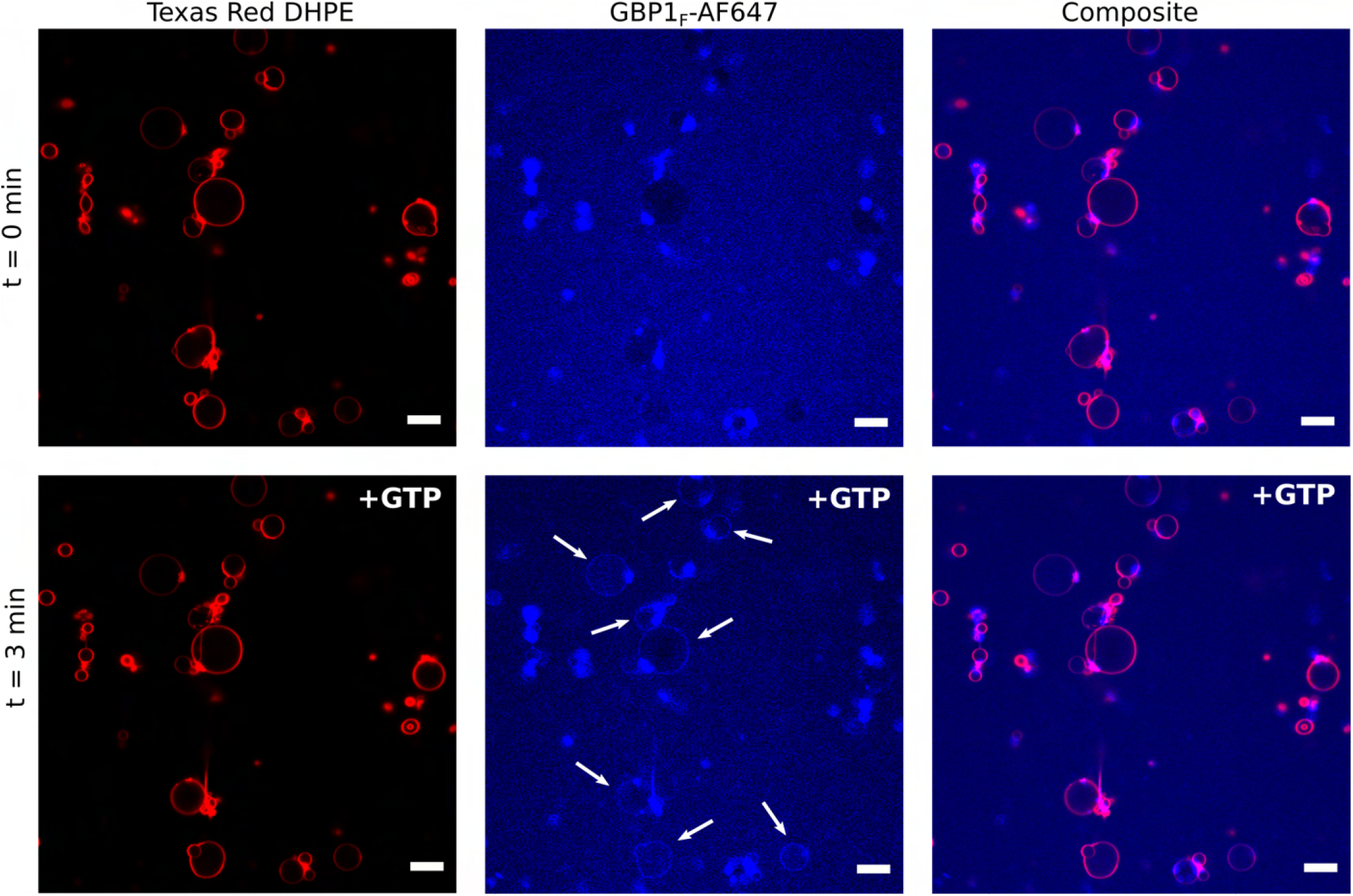
Confocal fluorescence imaging of GBP1*_F_* on GUVs. GBP1*_F_*-Q577C-AF647 shows weak binding to texas red-DHPE labelled GUVs 3 min after the addition of 1 mM GTP (white arrows). Scale bars correspond to 5 µm.

**Extended Data Fig. S15.**
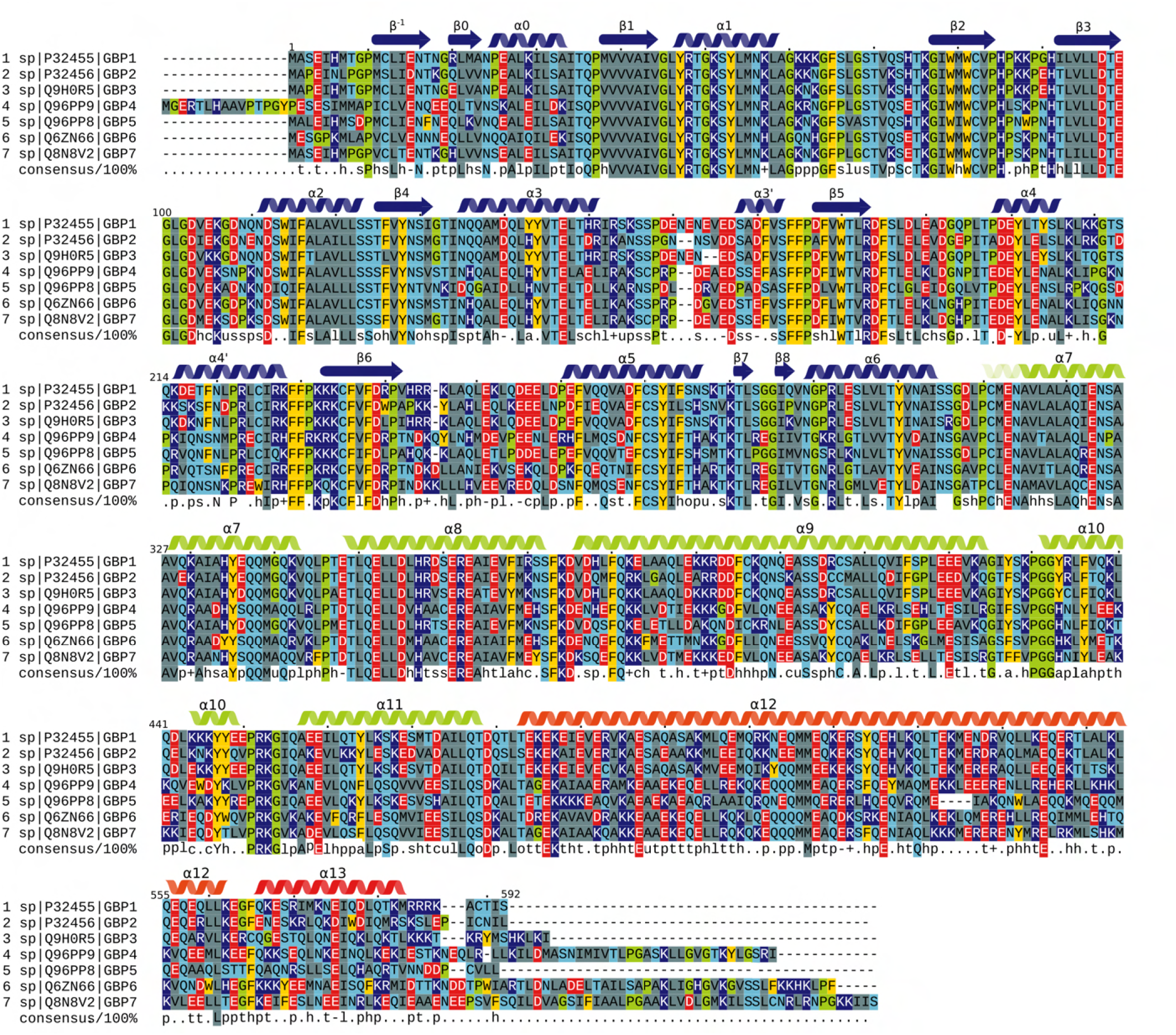
Multiple sequence alignment (MSA) of human GBP1-GBP7. Primary sequences of GBP1 - GBP7 (UniProt: P32455, P32456, Q9H0R5, Q96PP9, Q96PP8, Q6ZN66, Q8N8V2) were used as input for Clustal Omega (69). Secondary structure elements for GBP1 are displayed for guidance. The colour of the alpha-helices and beta-sheets correspond to the domain architecture of GBP1 shown in Fig. 1a and used throughout the main text (blue: Large GTPase domain, green: Middle domain (MD), orange/red: GTPase effector domain). Residues are coloured by physicochemical property of the side chain (grey: hydrophobic, light blue: polar, red: negatively charged, dark blue: positively charged, yellow: aromatic, green: special cases). The consensus sequence (100 %) is shown below the alignment together with conserved physicochemical classes (l: aliphatic, a: aromatic, c: charged, h: hydrophobic, -: negative, p: polar, +: positive, s: small, u: tiny, t: turnlike).

**Extended Data Fig. S16.**
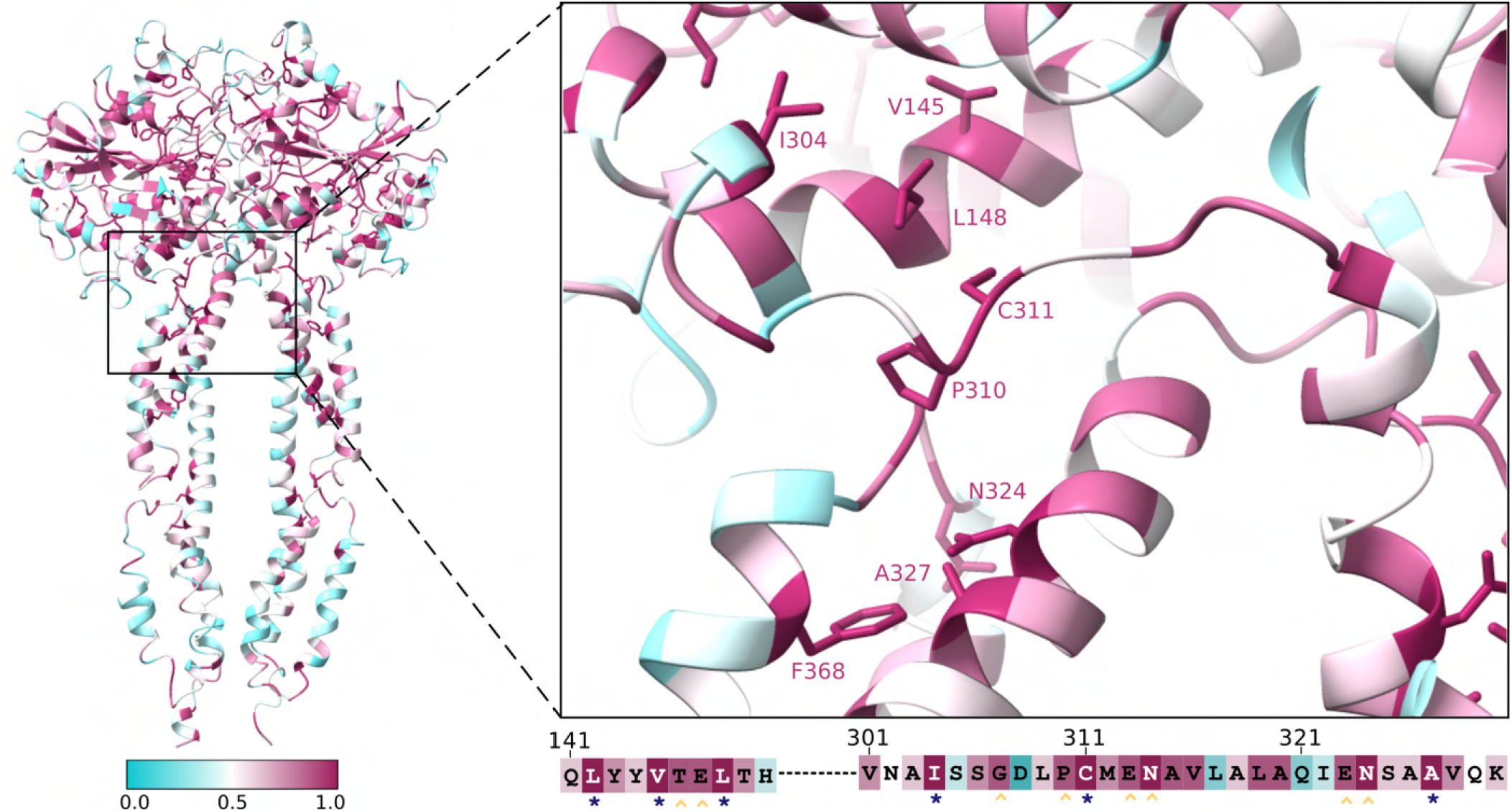
Sequence conservation mapped onto the structure of the GBP1-GDP*·*AlF_3_-dimer. Highly conserved regions are displayed in magenta whereas less conserved areas are shown in cyan. Residues with a conservation higher than 95 % are shown in stick representation. Right panel: Close-up of the cross-over linker region. The primary sequence of highly conserved stretches is displayed below (blue asterisk: highly conserved and buried residue, yellow circumflex: highly conserved and exposed residue).

**Extended Data Fig. S17.**
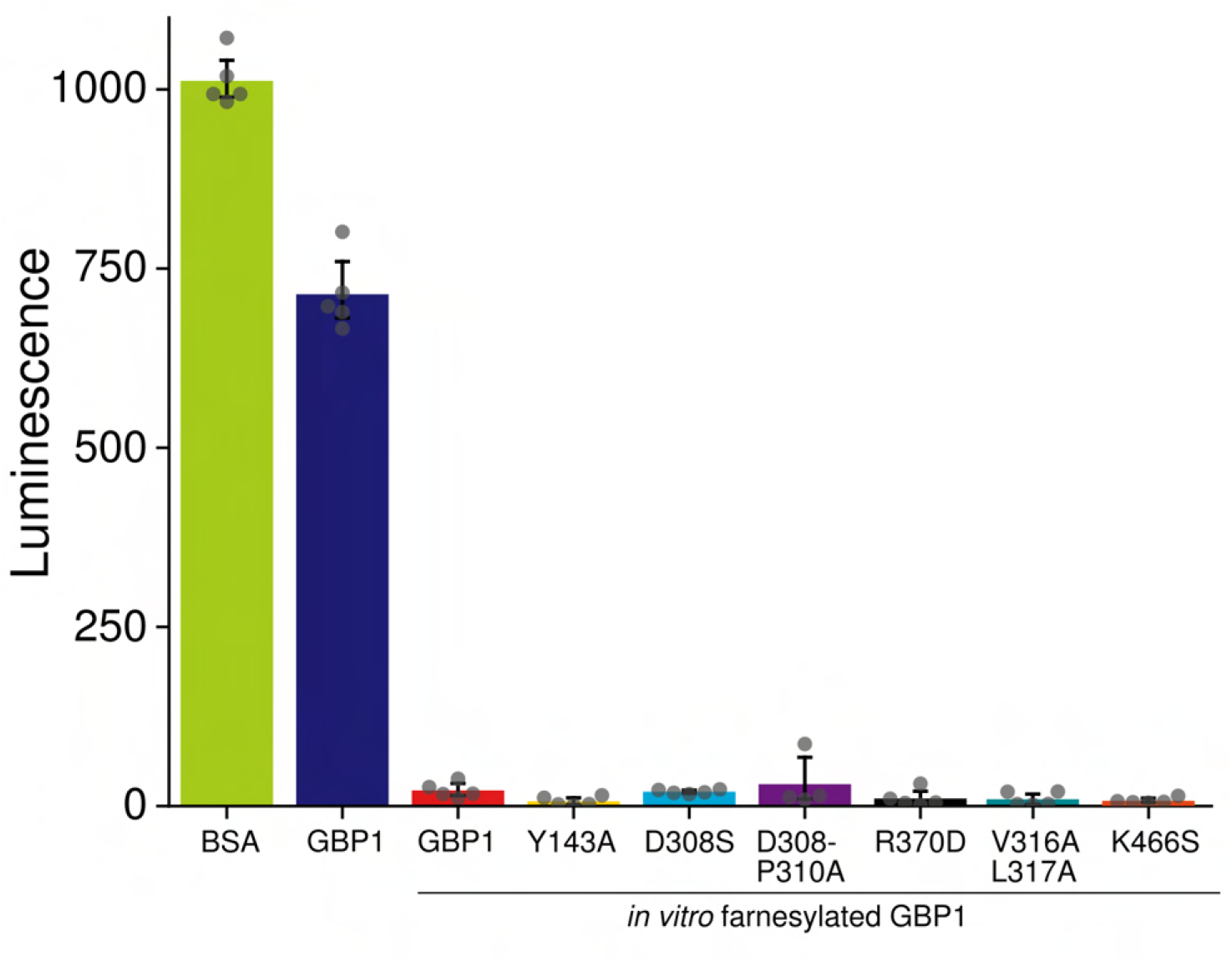
GTPase activity assay of GBP1. GTPase activity of non-farnesylated GBP1 and in vitro farnesylated GBP1 WT and GBP1 variants was determined using the GTPase-Glo™ Assay (Promega). Low luminescence signal corresponds to high GTPase activity. Bovine serum albumin (BSA) was used as a negative control (n=5).

**Extended Data Fig. S18.**
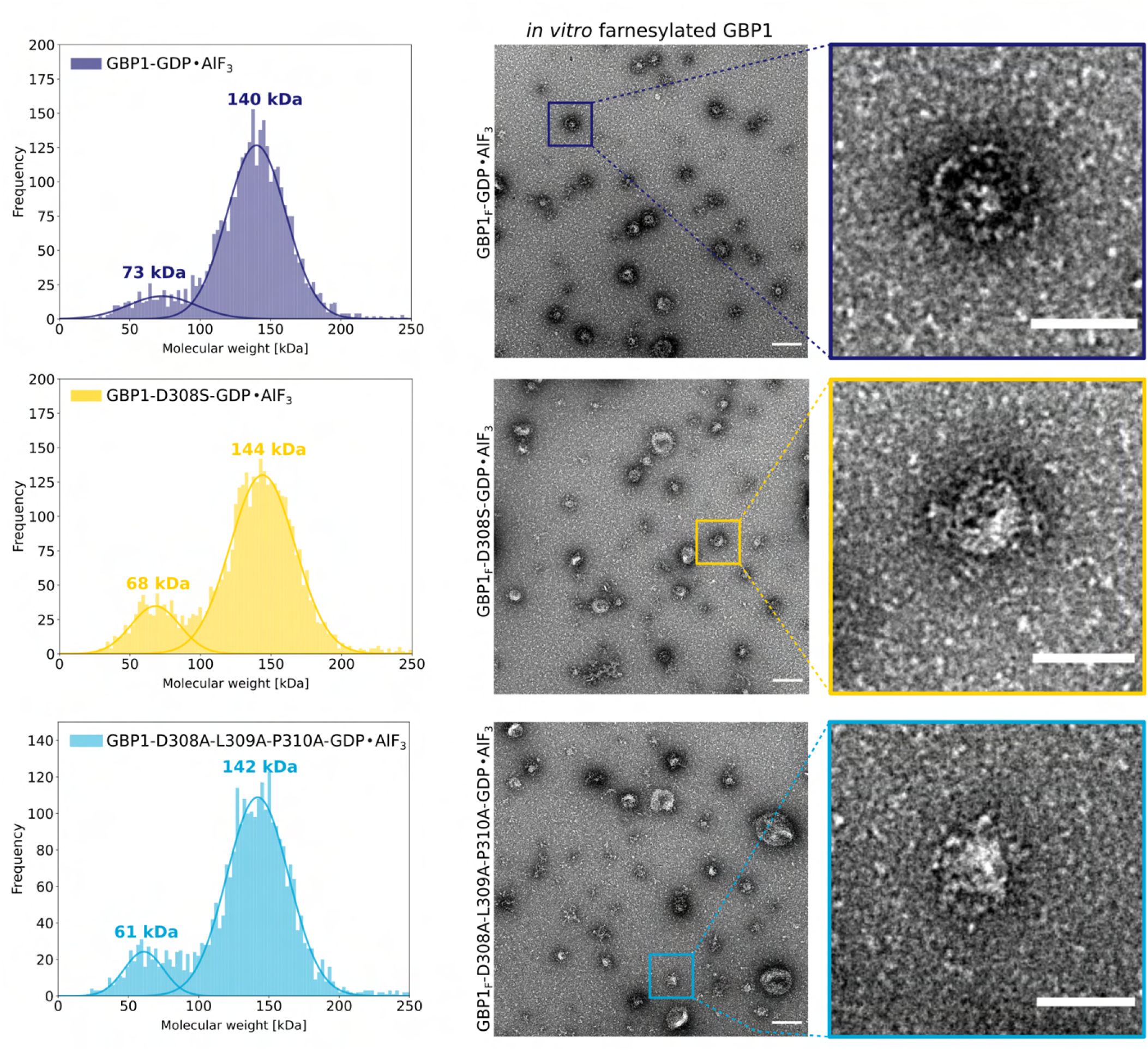
Effect of GBP1 variants on dimerisation and of GBP1*_F_* variants on membrane binding. Left panel: Individual mass photometry spectra revealing that variants D308S and D308A-L309A-P310A do not influence the ability of GBP1 to form dimers. Right panel: Negative stain EM of GBP1*_F_*, GBP1*_F_*-D308S and GBP1*_F_*-D308A-L309A-P310A bound to SUVs. A reduction in membrane binding for GBP1*_F_*-D308S and GBP1*_F_*-D308A-L309A-P310A is observed, but the capability to bind to membranes is not entirely lost. Scale bars correspond to 200 nm in the left column and 100 nm in the right column.

**Extended Data Fig. S19.**
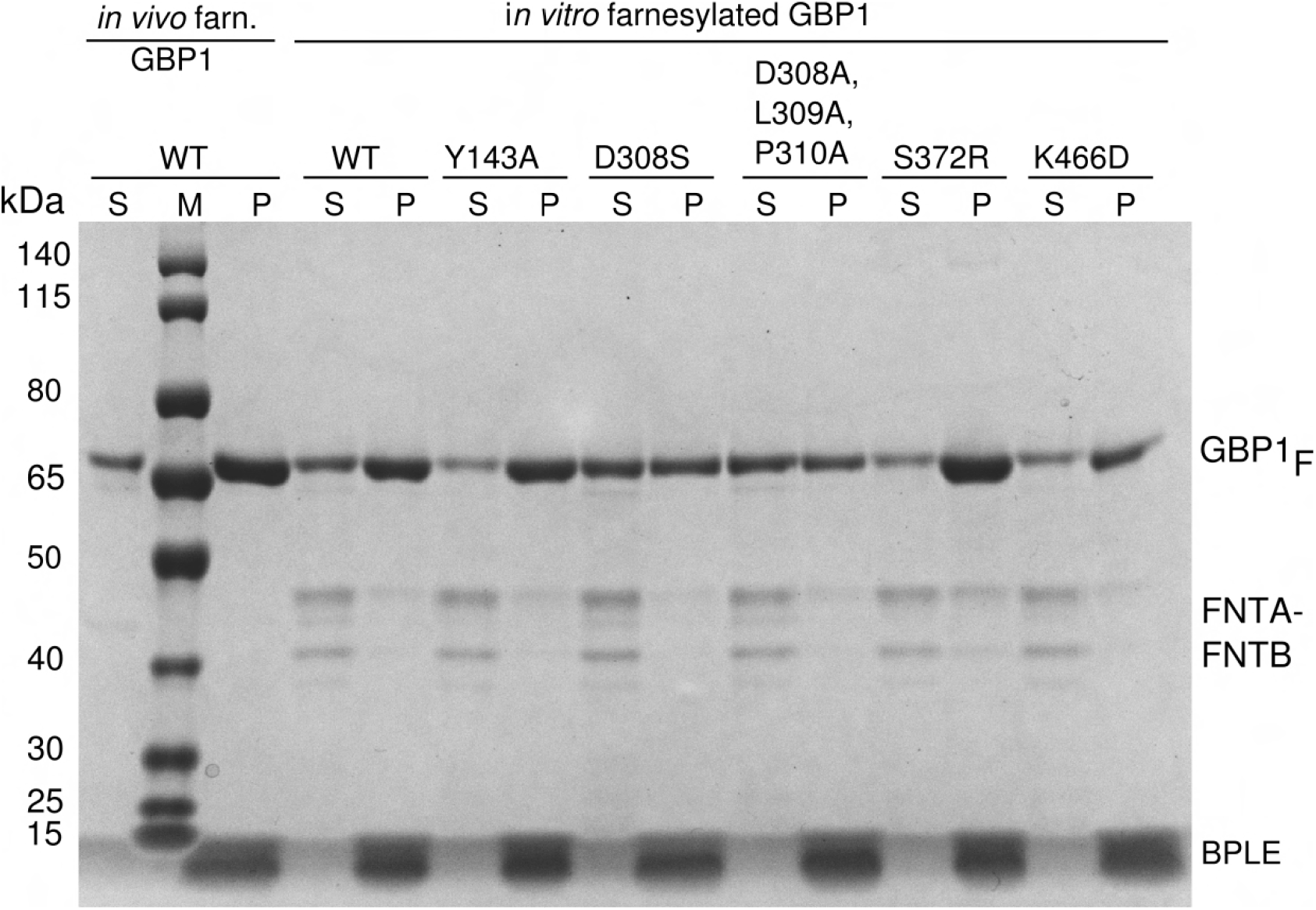
SDS-PAGE analysis of co-sedimentation assays. Representative uncropped SDS-PAGE gel from Fig. 6 showing the result of a co-sedimentation assay of GBP1*_F_* variants with BPLE-derived liposomes liposomes. S: Supernatant, P: Pellet

**Extended Data Fig. S20.**
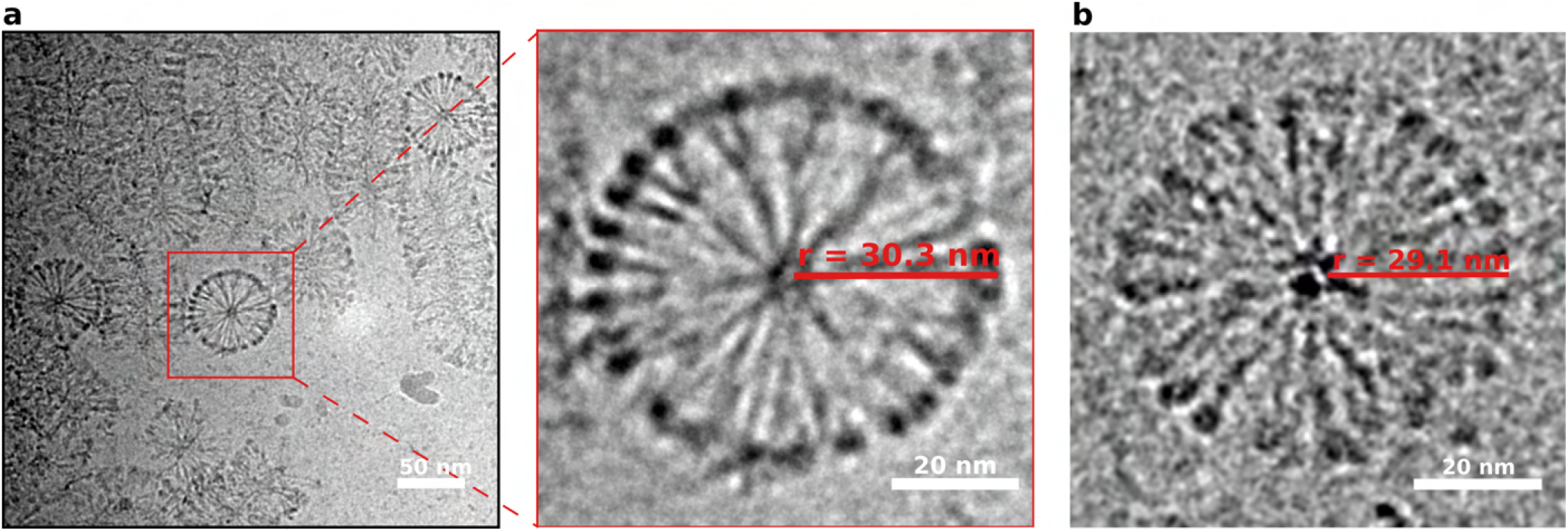
Top views of filaments formed by GBP1*_F_* on LPS-EB. (a) Putative top views of GBP1*_F_*-decorated LPS micelles are indicative of a highly ordered coat. The diameter of these projections was determined to be 60 nm. (b) For comparison, a GBP1*_F_* micelle with a diameter of 58 nm is shown.

**Extended Data Fig. S21.**
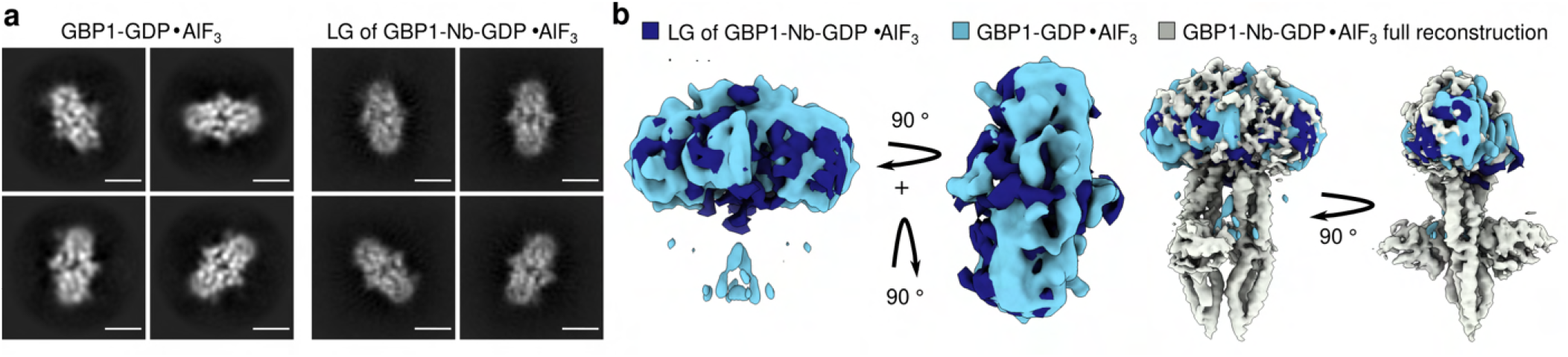
Comparison of 3D reconstructions of the LG dimer-like particles from the GBP1-GDP*·*AlF_3_ and GBP1-GDP*·*AlF_3_-Nb datasets. (a) 2D classes representative for projections of the LG domains of the GBP1 dimer from both datasets. Scale bar corresponds to 4 nm. (b) 3D reconstructions based on particles from 2D classes in (a) displaying density consistent with a dimeric LG domain. Rightmost panels: Comparison of both maps to the 3D reconstruction of GBP1-GDP*·*AlF3 -Nb dataset using all particles.

**Extended Data Table S1.** Summary of SEC-MALS measurements.

|  | Average molecular weight [kDa] | Number of measurements |
| --- | --- | --- |
| **GBP1** | 66.4 ± 2.1 kDa | n = 17 |
| **GBP1-GDP·AIF<sub>3</sub>** | 134.6 kDa ± 4.1 kDa | n = 16 |
| **GBP1<sub>F</sub>** | 66.1 kDa ± 0.3 kDa | n = 7 |
| **GBP1<sub>F</sub>-GDP·AIF<sub>3</sub>** | 67.6 kDa ± 1.7 kDa | n = 8 |
| **GBP1-Nb74** | 79.9 kDa ± 3.0 kDa | n = 3 |
| **GBP1-GDP·AIF<sub>3</sub>-Nb74** | 154.8 kDa ± 4.8 kDa | n = 6 |
| **GBP1<sub>F</sub>-Nb74** | 81.9 kDa | n = 1 |
| **GBP1<sub>F</sub>-GDP·AIF<sub>3</sub>-Nb74** | 81.9 kDa ± 3.5 kDa | n = 3 |

**Extended Data Table S2.** Summary of data collection and model refinement/validation statistics

|  | GBP1-GDP·AIF<sub>3</sub> | GBP1-GDP·AIF<sub>3</sub>-Nb74<br>EMD-16794<br>EMPIAR-11459 | GBP1<sub>F</sub>-GDP·AIF<sub>3</sub> |
| --- | --- | --- | --- |
| Data collection |  |  |  |
| Microscope | TFS Titan Krios | TFS Titan Krios | JEOL JEM3200FSC |
| Voltage (kV) | 300 | 300 | 300 |
| Detector | Quantum-K2 | Quantum-K3 | Quantum-K2 |
| Energy filter | Gatan Bioquantum | Gatan Bioquantum | In-column omega filter |
| Micrographs collected (no.) | 1193 | 5208 | 395 / 606 |
| Pixel size (Å) | 1.09 | 0.834 | 1.22 / 2.449 |
| Electron exposure (e-/Å<sup>2</sup>) | 60 | 60 | 48 / 13 |
| Frame number | 48 | 50 | 60 |
| Exposure time (s) | 10 | 2.2 | 12 |
| Defocus range (μm) | -0.5 to -3.5 | -0.6 to -2.2 | -1.0 to -3.0 |
| Data processing |  |  |  |
| Symmetry imposed | C2 | C1 or C2 |  |
| B-factor (Å<sup>2</sup>) | 231.1 | 158.3 (C1), 168.4 (C2) |  |
| Final number of particles | 35,715 | 181,161 |  |
| Final map resolution (Å) | 4.9 | 3.7 (C1), 3.6 (C2) |  |
| Map resolution range (Å) | * | 3.0 - 6.0 |  |
| Model refinement |  | PDB 8cqb |  |
| Initial model used (PDB code) |  | 1f5n and 2b92 |  |
| Primary sequence (Uniprot ID) |  | P32455 |  |
| Model resolution |  |  |  |
| FSC map-model (0.5)** |  | 3.97 (4.06 unmasked) |  |
| Map sharpening B-factor (Å<sup>2</sup>) |  | -158.3 |  |
| Model composition |  |  |  |
| Non-hydrogen atoms |  | 7688 |  |
| Protein residues |  | 956 |  |
| Nucleotides |  | 2 |  |
| Ligands |  | 4 |  |
| B-factors |  |  |  |
| Overall (Å<sup>2</sup>) |  | 74.77 |  |
| Protein (Å<sup>2</sup>) |  | 75.13 |  |
| Ligand (Å<sup>2</sup>) |  | 52.60 |  |
| Validation |  |  |  |
| MolProbity score |  | 0.94 |  |
| Clashscore |  | 5.28 |  |
| RMSD |  |  |  |
| Bond lengths (Å) |  | 0.005 |  |
| Bond angles (°) |  | 0.993 |  |
| Rotamer outliers (%) |  | 1.29 |  |
| Ramachandran plot |  |  |  |
| Favored (%) |  | 93.49 |  |
| Allowed (%) |  | 6.09 |  |
| Disallowed (%) |  | 0.42 |  |

\* Due to strong preferred orientation, the map resolution range for GBP1-GDP·AIF<sub>3</sub> is not conclusive.
\*\* Map-model FSC computed against globally sharpened map

**Extended Data Table S3.** Summary of data collection parameters of cryo-electron tomograms

|  | Tomogram 33 and Tomogram 39 | Tomogram 50 |
| --- | --- | --- |
|  | EMD-16813 and EMD-16814 | EMD-16815 |
| Content | GBP1<sub>F</sub>-GDP·AIF<sub>3</sub> + BPLE liposomes | GBP1<sub>F</sub>-GDP·AIF<sub>3</sub> + BPLE liposomes |
| Pixel size (Å) | 3.075 | 3.075 |
| Electron exposure (e-/Å<sup>2</sup>) | 93.94 (1.54 per increment) | 100.04 (1.64 per increment) |
| Tilt range (°) | -60 to 60 | -60 to 60 |
| Increment (°) | 2 | 2 |
| Acquisition scheme | Bidirectional | Bidirectional |
| Frame number | 10 per increment | 20 per increment |
| Exposure time per increment (s) | 2.0 | 3.0 |
| Defocus (µm) | -5.0 | -4.0 |

**Extended Data Table S4.** Summary of data collection parameters of LPS-datasets

|  | LPS-EB | LPS-EB-GBP1-GDP·AIF<sub>3</sub> | LPS-SM | LPS-SM-GBP1<sub>F</sub>-<br>GDP·AIF<sub>3</sub> | LPS-ST | LPS-ST-GBP1<sub>F</sub>-<br>GDP·AIF<sub>3</sub> |
| --- | --- | --- | --- | --- | --- | --- |
| Micrographs collected (no.) | 488 | 561 | 118 | 515 | 79 | 437 |
| Pixel size (Å) | 1.288 | 1.288 | 2.449 | 1.288 | 1.891 | 1.891 |
| Electron exposure (e-/Å<sup>2</sup>) | 46.68 | 48.01 | 16.15 | 43.01 | 32.02 | 26.53 |
| Frame number | 50 | 50 | 50 | 50 | 60 | 48 |
| Exposure time (s) | 15 | 15 | 15 | 15 | 15 | 12 |
| Defocus range (μm) | -1.0 to -<br>3.5 | -1.0 to -3.0 | -1.0 to -<br>3.5 | -1.0 to -3.5 | -1.0 to -<br>3.5 | -1.0 to -3.5 |

**Extended Data Table S5.**
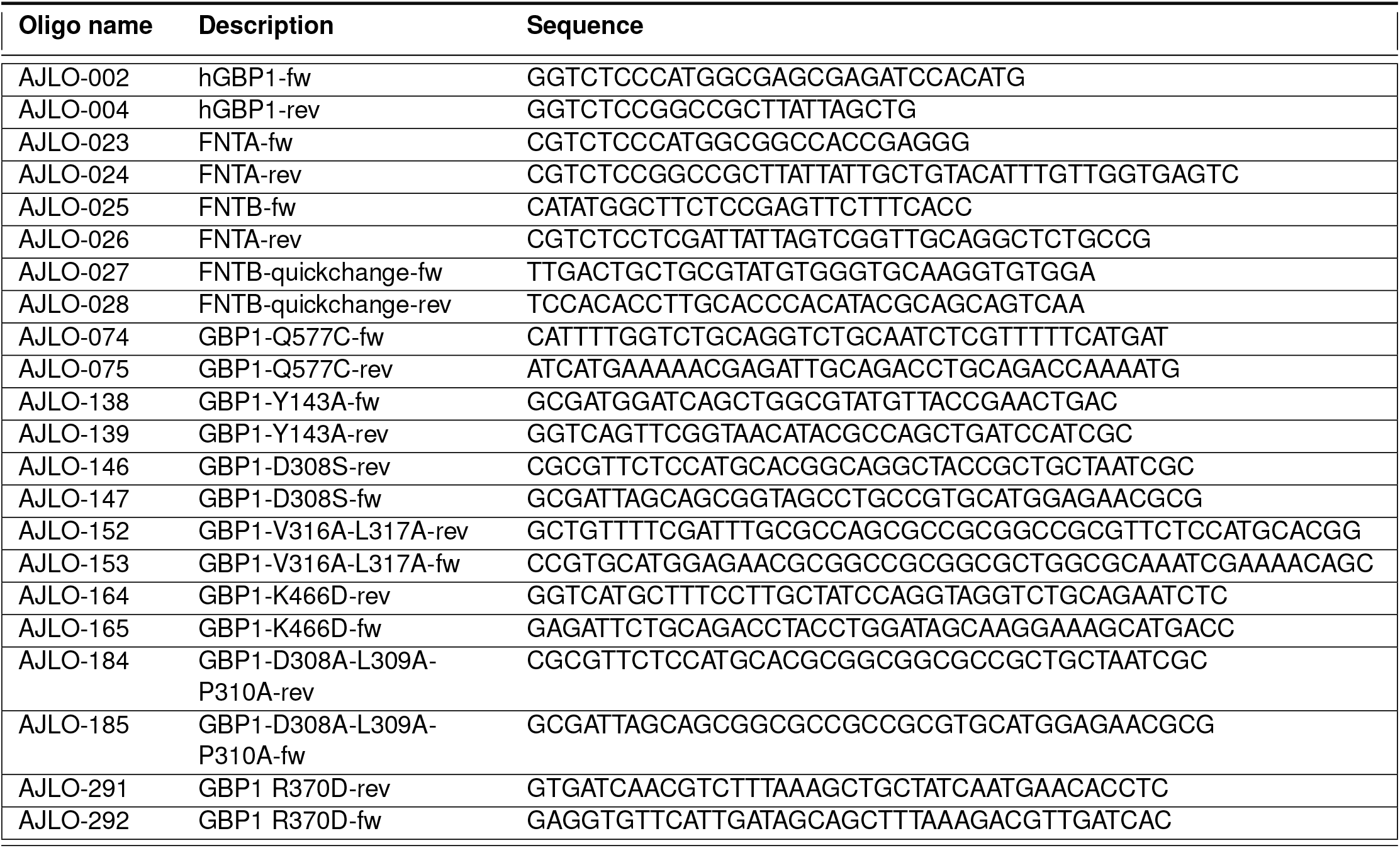
Primer sequences used in this manuscript.

**Extended Data Table S6.** Vectors and Constructs used in this manuscript.

| Name | Description | Source/ Reference |
| --- | --- | --- |
| AJLV0009 | pETM14: *E.coli* expression vector (KanR) | (73) |
| AJLV0038 | pCDFDuet: *E.coli* expression vector (SmR) | Merck Millipore (Novagen) |
| AJLV0040 | pJET1.2/blunt: Positive selection cloning vector (AmpR) | Thermo Fisher Scientific |
| AJLD0001 | Subcloning vector pUC57 with synthetic gene of hGBP1 optimised for *E.coli* expression (AmpR) | GenScript |
| AJLD0007 | GST-tagged-FNTA *in vitro* expression vector pANT7 (AmpR) | DNASU clone HsCD00630808 |
| AJLD0008 | GST-tagged-FNTB *in vitro* expression vector pANT7 (AmpR) | DNASU clone HsCD00733069 |
| AJLD0022 | NdeI site removed from pANT7-FNTB-cGST (AmpR) | derived from AJLD0008 |
| AJLD0030 | pETM14-hGBP1: *E.coli* expression vector (KanR) |  |
| AJLD0031 | pCDF-Duet-FNTA: Intermediate vector for cloning purposes (SmR) | Derived from AJLV0038 and AJLD0007 |
| AJLD0035 | pCDFDuet-FNTA-FNTB: *E.coli* co-expression vector for *in vivo* farnesylation of hGBP1 (SmR) | Derived from AJLV0038 and AJLD0022 |
| AJLD0052 | pJET1.2 His6-FNTA: Intermediate vector for cloning purposes (AmpR) | Derived from AJLV0040 and AJLD0007 |
| AJLD0053 | pJET1.2 FNTB (NdeI removed): Intermediate vector for cloning purposes (AmpR) | Derived from AJLV0040 and AJLD0022 |
| AJLD0056 | pETM14-hGBP1-Q577C: *E.coli* expression vector (KanR), point mutation for site-specific labelling | Derived from AJLD0030 |
| AJLD0063 | pCDFDuet-His-FNTA-FNTB: *E.coli* co-expression vector (SmR) to express His-FNTA-FNTB | Derived from AJLV0038, AJLD0052 and AJLD0053 |
| AJLD0074 | pMES4y-CA16697: C-His6-tagged nanobody (Nb74) raised against farnesylated GBP1 | Instruct-ERIC (PID7267) |
| AJLD0147 | pETM14-hGBP1-Y143A: *E.coli* expression vector (KanR) with point mutation | Derived from AJLD0030 |
| AJLD0150 | pETM14-hGBP1-L316A-V317A: *E.coli* expression vector (KanR) with point mutation | Derived from AJLD0030 |
| AJLD0151 | pETM14-hGBP1-D308S: *E.coli* expression vector (KanR) with point mutation | Derived from AJLD0030 |
| AJLD0153 | pETM14-hGBP1-D308A-L309A-P310A: *E.coli* expression vector (KanR) with point mutation | Derived from AJLD0030 |
| AJLD0158 | pETM14-hGBP1-K466D: *E.coli* expression vector (KanR) with point mutation | derived from AJLD0030 |
| AJLD0244 | pUC57-hGBP1-R370D: Subcloning vector with point mutation | Derived from AJLD0001 |
| AJLD0247 | pETM14-hGBP1-R370D: *E.coli* expression vector (KanR) with point mutation | Derived from AJLD0244 and AJLV0009 |

**Extended Data Table S7.** Bacterial host strains used in this manuscript.

| Strain | Description | Reference |
| --- | --- | --- |
| *E.coli* DH5α | Cloning host | Invitrogen |
| *E.coli* BL21(DE3) | Expression host | Thermo Fisher Scientific (74) |
| *E.coli* WK6 | Expression host for Nanobodies | Zell & Fritz, 1987 (75) |

