## Supplementary figures and images for "Structural basis of membrane targeting and coatomer assembly by human GBP1"

### Movie SM2

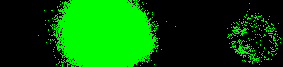
